# High-resolution structures of the UapA purine transporter reveal unprecedented aspects of the elevator-type transport mechanism

**DOI:** 10.1101/2024.08.23.609436

**Authors:** George Broutzakis, Yiannis Pyrris, Ifigeneia Akrani, Alexander Neuhaus, Emmanuel Mikros, George Diallinas, Christos Gatsogiannis

## Abstract

UapA is an extensively studied elevator-type purine transporter from the model fungus *Aspergillus nidulans*. Determination of a 3.6Å inward-facing crystal structure lacking the cytoplasmic N-and C-tails, molecular dynamics (MD), and functional studies have led to speculative models of its transport mechanism and determination of substrate specificity. Here, we report full-length cryo-EM structures of UapA in new inward-facing apo- and substrate-loaded conformations at 2.05-3.5 Å in detergent and lipid nanodiscs. The structures reveal in an unprecedented level of detail the role of water molecules and lipids in substrate binding, specificity, dimerization, and activity, rationalizing accumulated functional data. Unexpectedly, the N-tail is structured and interacts with both the core and scaffold domains. This finding, combined with mutational and functional studies and MD, points out how N-tail interactions couple proper subcellular trafficking and transport activity by wrapping UapA in a conformation necessary for ER-exit and but also critical for elevator-type conformational changes associated with substrate translocation once UapA has integrated into the plasma membrane. Our study provides detailed insights into important aspects of the elevator-type transport mechanism and opens novel issues on how the evolution of extended cytosolic tails in eukaryotic transporters, apparently needed for subcellular trafficking, might have been integrated into the transport mechanism.

## Introduction

Despite evolutionary, structural, and functional differences, elevator-type transporters use a common mechanism to translocate nutrients and metabolites across cell membranes. A mobile core domain (the *elevator*), including a single substrate binding site, slides along a more rigid scaffold domain^1–3^ to release the substrate on the opposite side of the membrane. In most cases, the scaffold serves as a dimerization or trimerization domain that supports the sliding of the core, a challenging process that dynamically modifies interactions with membrane lipids^4–7^. Substrate binding, translocation, and release elicit local structural changes necessary for the reversible sliding of the core and characteristic distinct transporter conformations^2^.

UapA, the uric acid-xanthine/H^+^ symporter in the model filamentous fungus *Aspergillus nidulans,*^8^ has historically defined the ubiquitous Nucleobase Ascorbate Transporter (NAT) family^9^ (also known as SLC23^10^). NAT transporters are specific for purines, pyrimidines, and related metabolites (e.g., nucleosides, cytokines) or analogs used as first-line drugs (e.g., 5-fluorouracil, allopurinol, allopurinol, thiopurines)^11–14^. Surprisingly, the subgroup of mammalian NATs characterized functionally is highly specific for L-ascorbate/Na^+^ symport (SVCT1/2). Ascorbate-specific NATs, essential proteins in mammals and plants^15^, appear to have evolved in vertebrates. Interestingly, anthropoid primates have lost the group of nucleobase-specific NATs, retaining only ascorbate-specific members^16^.

The plethora of classical and reverse genetic tools, coupled with functional and *in vivo* cellular biology studies possible in *A. nidulans,* have made UapA a model transporter to address practically all aspects of elevator-type transport mechanism and physiological function^14,17,18^. The elucidation of the UapA crystal structure in 2016 in the xanthine-bound cytoplasm-facing inward-open state, despite its intermediate resolution at 3.6Å, was invaluable in confirming the findings of past and ongoing functional and genetic studies regarding the localization of specificity-modifying residues and the importance of dimerization for transporter activity^4^.

For other NATs, there are, at present, only a handful of structures available. These correspond to two distinct topologies of the bacterial uracil permease UraA^19,20^, the mouse and human SVCT1 ascorbate transporters^21,22^, and the plant Azg1 cytokine transporter^23^, all corresponding to cytoplasm-facing topologies, albeit displaying apparent structural differences. In the case of hSVCT1, there is also a distinct topology in which the substrate binding cavity is occluded from both the cytoplasm and the exterior^22^. Few structural studies on similar transporters have successfully trapped outward-facing (i.e., extracellular-facing) or additional states, but these transporters are phylogenetically and functionally very distinct from NATs, as is the case of the human erythrocyte AE1/Band3 Cl^-^/HCO3^-^ anion exchanger 1^24–26^ and the boron transporter Bor1^27,28^ of the SLC4 family, or the nucleoside CNT3 transporter of the SLC28 family^29^.

Notably, a largely ignored aspect of transporter structure analysis is the specific roles of eukaryotic cytoplasmic N- or C-tails, most often missing from prokaryotic homologs. Cytoplasmic tails have been shown to play prominent roles in the folding, biogenesis, and turnover of eukaryotic transporters, transport kinetics, and, surprisingly, substrate specificity^30–32^. For UapA specifically, the N-tail has been shown to affect proper folding, ER-exit, trafficking to the PM, and possibly dimerization^33,34^. Interestingly, the closest prokaryotic UapA homologs lack such N-tail elements^35^. For example, the *Escherichia coli* homologs XanQ^36^ and UgfU^37^, which possess highly similar kinetic characteristics and specificity to UapA (xanthine and uric acid transporters), have much shorter and distinct N-tails. The UraA uracil transporter^20^ has an even shorter cytosolic N-terminal of only 12 amino acids. The C-tail of UapA has an essential role only in the endocytic turnover of the transporter in response to stress signals^38,39^.

In this study, we conducted a comprehensive cryoEM analysis, yielding several structures of nearly full-length wild-type (WT) UapA in its apo and substrate-bound states in detergent. We also resolved the cryo-sensitive and altered specificity mutant Q408E^34^ in detergent and lipid nanodiscs. The structures reveal new inward-open conformations, provide the structural basis of substrate binding and specificity, and rationalize functional studies accumulated over the last decades. The 2.06Å resolution reported here for the apo state of UapA in detergent stands to date as the highest ever achieved for a eukaryotic transporter using any structural method, allowing us to elucidate the critical role of solvent molecules in ligand binding and the network of functionally important protein-lipid interactions. Notably, in all high-resolution structures, the cytoplasmic N-tail domain is structured. It shows multiple functionally important interactions with both the core and scaffold domains, opening new issues regarding a regulatory role in transport activity. The structural results, along with functional studies and MD simulations, dissect UapA architecture and the molecular basis of substrate-specific recognition with unmatched detail. They offer insights into previously elusive aspects of the transport mechanism, including the role of lipids and water molecules in specificity, dimerization, and transport, and highlight the significant functional roles that cytosolic tails may have in eukaryotic elevator-type transporters.

## Results

### The overall architecture of UapA

We expressed UapA and a set of different variants in *Saccharomyces cerevisiae*^40^. All constructs were truncated for the first 10 N-terminal residues, a deletion shown to increase protein stability and be advantageous for protein purification while maintaining WT activity^4^ and contained a C-terminal GFP-His_8_ tag, which does not affect the kinetics of UapA-mediated transport^9,41^. For the rest of this article, we will refer to UapA_Δ1-10_GFP_8xHis_ as UapA_WT_. Purified UapA_WT_ in detergent (DDM) with and without xanthine was subjected to cryoEM, and we obtained inward-open structures in both the apo- and the substrate-bound state, at an average resolution of 2.06 and 2.4 Å, respectively (**Supplementary Figures 1** and **2**; UapA_WT_-Apo-DDM, UapA_WT_-Xan- DDM, respectively). Furthermore, we obtained the inward-open structure of the cryo-sensitive variant Q408E in the apo state in detergent at an average resolution of 2.6Å (**Supplementary Figure 3**; UapA_Q408E_-Apo-DDM). Finally, to obtain detailed insights into UapA-lipid interactions, we reconstituted UapA_Q408E_ in nanodiscs and obtained a 3.5 Å structure of the substrate-bound state in a close-to-native lipid environment (**Supplementary Figure 4**; UapA_Q408E_-Xan-ND). The four structures display distinct inward-open (hereafter called IO) conformations and show characteristic intriguing structural differences that are not only restricted to substrate-binding, which will be discussed and highlighted in detail in the following sections. For clarity, we first describe the overall IF UapA architecture based on the best-resolved cryoEM structure of this study, the 2.06 Å UapA_WT_-Apo-DDM.

UapA forms a homodimer with a total molecular weight of ∼120 kDa. Dimer assembly is essential for UapA function as disruption of intra-protomeric interactions drastically reduces activity^33^. Furthermore, lipids that co-purify with UapA are indispensable for dimer formation and, thus, for activity^6^. In the novel cryoEM structure, each protomer is organized in three domains: the N-tail, the core (elevator), and the scaffold (dimerization) domain. Similar to all known NATs, UapA consists of 14 transmembrane segments (TMS) arranged in a 7+7 inverted repeat architecture. TMS 1-4 and 8-11 constitute the core domain (**Figure 1**, blue segment), which contains the substrate binding site in a cavity delineated by the half helices of TMS3 on the extracellular side and TMS10 on the cytosolic side. TMS5-7 and 12-14 form the more stable scaffold domain (**Figure 1**, pink segment), which provides the surface against which the core domain slides when undergoing the inward-to-outward conformational shift but also provides the dimerization interface.

**Figure 1:**
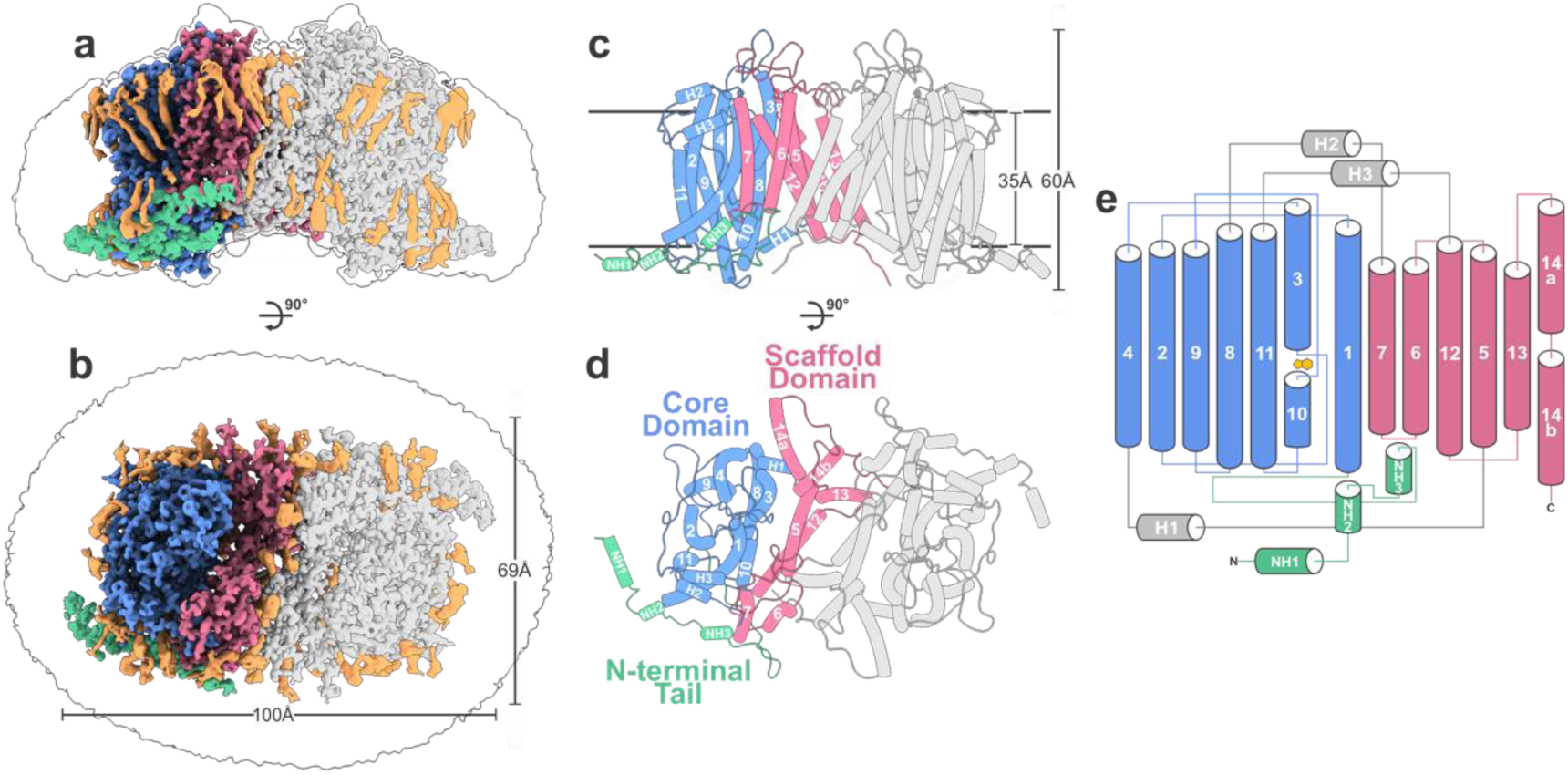
Structure of apo-UapA in detergent. **a-b)** cryo-EM structure of the UapA dimer (UapA_WT_-Apo-DDM) as viewed from the side (a) and from the extracellular space facing the cytoplasm (b). One protomer is colored by domain, with the core domain colored blue, the scaffold domain pink, and the N-terminal tail green. Densities assigned to lipid or detergent molecules are colored in orange. The outline marks the end of the detergent micelle. This coloring scheme is maintained throughout the paper. **c-d)** corresponding molecular models. **e)** Topology diagram of UapA. α-helices are represented by cylinders and loops by thin lines. Xanthine is drawn in yellow.

The cryoEM structure of UapA_WT_-Xan-DDM closely resembles the previous 3.6 Å crystal structure of IF substrate-bound UapA^4^ (RMSD 0.7Å) (**Supplementary Figure 5)** but reveals essential differences. Notably, the extracellularly facing loops connecting odd to even numbered TMS show different conformations compared to the crystal structure, with an overall RMSD of 1.027 Å (**Supplementary Figure 5a**). Furthermore, in addition to the already established and functionally important disulfide bridge in the loop following TMS3 (L3), we identified a second disulfide bridge on L5 (**Supplementary Figure 5b-c**). The difference in the conformation of this loop is not surprising, given that the region is involved in the crystallization interface in the previously reported crystal structure. The unprecedented level of detail of the cryoEM map allowed the modeling of ∼300 water molecules and 62 detergent tails and/or endogenous co-purified lipids. The intricate role of the water network and lipids in substrate binding and stabilization of the binding site, as well as the role of lipids in dimer formation and activity, will be discussed in a later section.

A striking finding across all cryoEM structures is that the N-tail (residues 28-74) is structured and well resolved, in contrast to the crystal structure. This region, which was long thought to be unstructured, has been shown to be essential for proper folding, ER exit, and trafficking but has not been anticipated to play a direct role in transport activity^33^. Our new structures reveal for the first time that the N-tail forms a characteristic helical lid, capping both the scaffold and the core domain within a monomer (**Figure 1**, green). This critical finding opens new issues on the evolution of extended cytosolic tails in eukaryotic transporters, as these are much shorter or missing from bacterial homologs that show very similar transport activity^30^.

### The UapA N-tail interacts functionally with both the scaffold and core domains

We have previously performed a semi-systematic analysis of the 74-amino-acid long N-tail of UapA by mutational analysis. Deleting the first 30 residues did not affect proper UapA biogenesis or UapA-mediated growth on uric acid or xanthine. In contrast, longer deletions led to partial or total ER retention^33^. The N-tail in the cryoEM structure of UapA_WT_-apo-DDM is resolved starting from residue Gln26, thus close to the position where the N-tail begins to include functionally essential elements.

The N-tail consists of three short alpha helices, termed NH1-3, followed by an extended loop, and it closely interacts with both the core and the scaffold domains (**Figure 1**; **Figure 2a**). NH1 exhibits a transverse orientation, lying on the surface of the membrane, as evidenced by its position relative to the rest of the protein and its largely amphipathic nature, hydrophobic proximally to the membrane while strongly positive distally (**Figure 1; Supplementary Figure 6a-d)**. MD further validated this positioning, showing NH1 buried amongst the head-groups of the phospholipid bilayer (**Supplementary Figure 6e)**. The rest of the tail follows the cytoplasmic edge of the transporter, connecting the core and scaffold domains by stably interacting with both (**Figure 2a**). The N-tail thus resembles a structural lid, displaying characteristic shape and charge complementarity with the body of the transporter in three distinct regions (**Figure 2b**).

**Figure 2:**
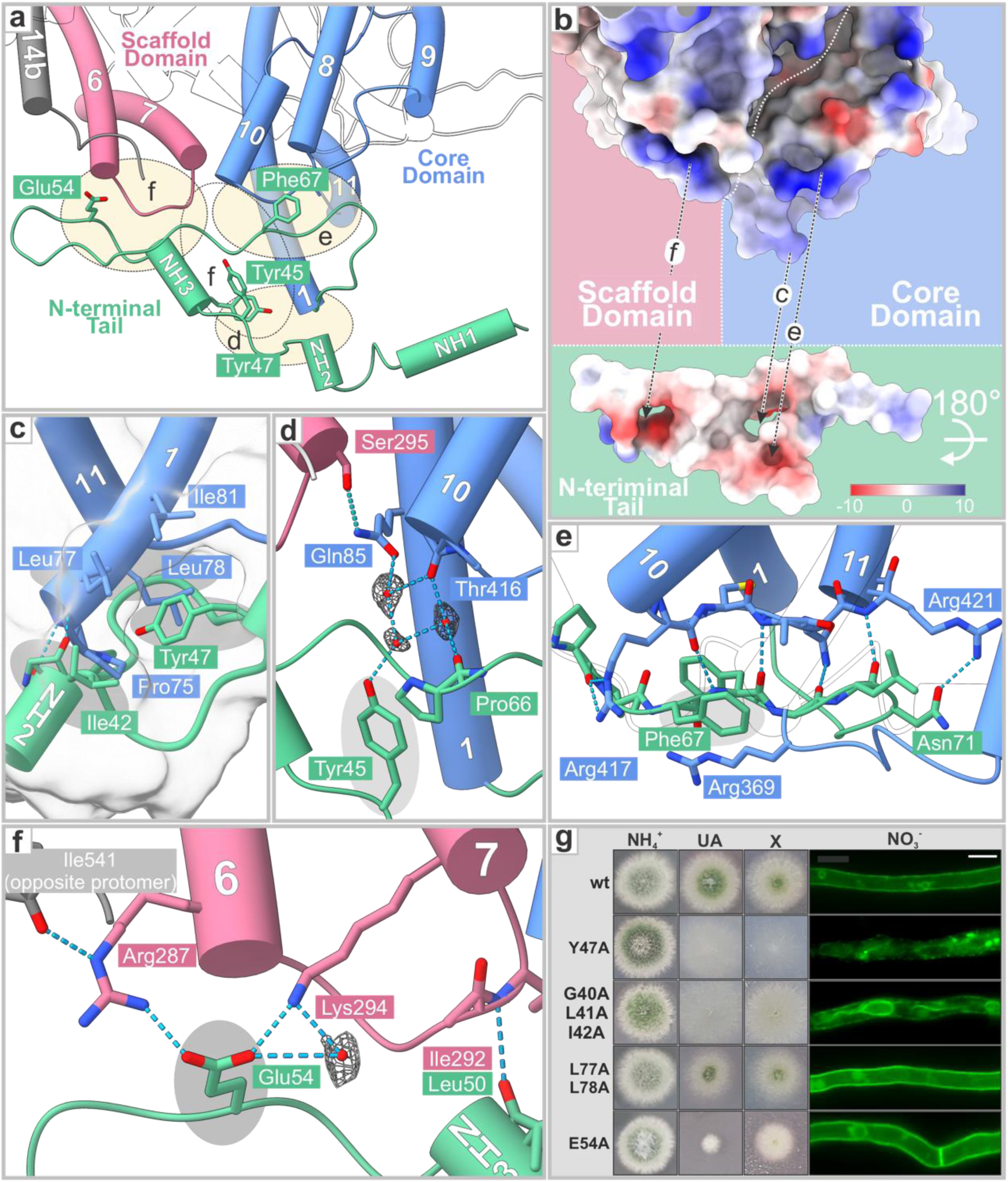
Inter-domain interactions of the N-terminal tail. **(a)** Overview of the N-tail interactions, as viewed from the cytoplasm. Selected important residues are shown. **(b)** Surface Coulombic potential of the interacting interphases. The N-terminal domain is pictured below, translated away from the main body of the transporter and rotated 180° about its long axis. The main body of the protein is viewed from the same angle as in (a). Units in kcal/(mole) at 298 K. **(c)**-**(e)** Interactions between the tail and the core domain. **(c)** Shape complementarity between Tyr47 and the hydrophobic surface created by Ile42 and TMS1. Surface representation of the region, excluding Tyr45, is shown in gray. **(d)** The water network surrounding Tyr45 is part of the conserved DYDY motif of the tail. **(e)** β-sheet-like interactions of Pro66-Asn71 of the tail with Arg417-Arg420 of L10 in the core domain. Notice the cation-π interaction of Phe67-Arg369 of L8. **(f)** The interaction between the tail and the scaffold domain is mainly facilitated by the double salt bridge between Glu54 of the tail and Arg287, Lys294. Hydrogen bonds are displayed as dashed, cyan lines, and the densities corresponding to water molecules are displayed as gray mesh **(g)** Functional analysis of selected N-terminal mutants. Notice the toxicity phenotype (i.e., reduced colony radius and green conidiospore production) in E54A and to a lower degree in L77A/L78A.

#### Interaction of the N-tail with the core domain

The N-tail closely interacts with the core domain through numerous hydrogen bonds on three main surfaces (**Figure 2c-e**). The first of those interfaces stabilizes the tail to TMS1. NH2 functions as a cytoplasmic extension of TMS1, with two α-helical-like hydrogen bonds stably linking the two segments. Directly medially to that interface, we observe a hydrophobic pattern complementarity between the first few residues of TMS1 with the short loop NL2 linking NH2 and NH3. More specifically, Tyr47 of NL2 snugly fits into a hydrophobic cavity formed by Ile42 in NH2 and Leu77, Leu78, and Ile81 in TMS1. This N-tail-TMS1 interaction is stabilized further by a hydrogen bond between the phenolic group of Tyr47 and the backbone =O of Ile42 (**Figure 2c**). Mutational analyses performed previously and herein strongly support that the residues involved in N-tail-TMS1 interactions are functionally important. Tyr47 mutations to non-aromatic residues lead to massive perinuclear ER retention, abolishing substrate-mediated growth while preserving a wild-type *K*_m_^33^ (**Figure 2g**), suggesting proper substrate binding site folding in the ER-retained variant.

Similarly, but to a lesser extent, specific mutations affecting Ile42, which forms the cytoplasmic barrier of the hydrophobic pocket, increase ER retention (**Figure 2g, Supplementary Figure 7**). A double Ala replacement of Leu77/Leu78 leads to elevated substrate transport, as suggested by increased uric acid toxicity in growth tests (i.e., reduced colony diameter; see **Figure 2g** and **Supplementary Figure 7**). Thus, the N-tail-TMS1 interaction could affect both proper ER exit and transport kinetics. All involved residues (Tyr47, Ile42, Leu77) are well conserved in the NAT family.

The high resolution achieved in this study also allowed us to identify a water network between Tyr45 and Pro66 of the N-tail and Thr416 and Gln85 of the core domain (**Figure 2d**), unveiling a second N-tail-core interface. Mutating Tyr45 alone leads to a very moderate effect in trafficking, but co-mutating its neighboring region (Asp44, Asp46, and Tyr47) results in a much more extreme trafficking defect than mutating Tyr47 alone^33^. It is also interesting to notice that Gln85 is positioned close to the binding site and functionally links the core and the scaffold domain via its interaction with Ser295 of the scaffold domain (**Figure 2d**). Thus, Gln85 is a residue that directly links the three domains of the transporter, which rationalizes and underlines its absolute conservation in the entire NAT family^4,41,42^.

Finally, the third and most extensive interface includes NL3, the long loop following NH3, which runs parallel with the cytosolic loop L10. Here, the extended sidechain and backbone-level β-sheet-like interaction, along with the very strong and characteristic cation-π bond between Phe67 in NH3 and Arg369 in the proximal L8, makes significant conformational changes in the region during the translocation cycle less likely (**Figure 2e)**. Note that a P66A/F67A mutant is partially ER retained, supporting the functional importance of these residues^33^ (**Supplementary Figure 7**).

#### Interaction of the N-tail with the scaffold domain

While the N-tail is firmly bound to the core domain via distinct and extended interfaces, the interaction with the scaffold is established by a singular bidentate salt bridge (**Figure 2a, f**). Glu54, situated on NL3, interacts with Lys294 and Arg287 on cytosolic loop L6, further stabilizing the former by a water molecule (**Figure 2f)**. Peculiarly, Glu54 is not a conserved residue in homologs of UapA, while its proposed interacting partners, Lys294 and Arg287, are very well conserved. An intriguing finding is that mutating Glu54 leads to characteristic uric acid toxicity, as evident from growth tests, while proper trafficking to the plasma membrane remains unaffected (**Figure 2g**). Previously identified phenotypically equivalent mutations have been identified in TMS12 on the core/scaffold interface directly opposite the substrate binding site^43^. Conceptually, uncoupling the N-tail from the scaffold domain (Glu54 mutation) and increasing the sliding between the core and the scaffold (TMS12 mutations) are similar in that they decrease the energy required for the conformational shift, increasing the speed and thus the capacity of the transporter to such an extent that the substrate becomes toxic. These findings strongly suggest that the N-tail-core-scaffold interactions, besides their absolute essentiality for ER exit and proper trafficking, are also crucial for controlling the transport dynamics of UapA *per se*.

### Molecular Dynamics suggest that the N-tail remains bound to the core and scaffold domains during the sliding of the elevator domain and transport catalysis

A critical emerging question is whether the characteristic N-tail lid, which caps both the core and scaffold domains, permits sliding of the core domain against the scaffold during the translocation cycle or if it instead locks the transporter in an IF state and must be disengaged before transitioning to the occluded (Occ) and outward-facing (OF) state. The high-resolution structures for xanthine-bound and apo-IO UapA allowed us to construct improved and confident homology models for the ‘missing’ Occ und OF conformations of UapA to address such questions and gain insights into the translocation mechanism and elusive structure-function relationships. These models are based on the structures of UraA (PDB: 5XLS) for the Occ state and human AE1 (PDB: 4YZF) for the OF state. MD simulations of the homology models suggest that the N-tail remains bound to the core and scaffold domains throughout the translocation cycle but extends to accommodate the sliding of the core domain against the scaffold **(Figure 3)**.

**Figure 3:**
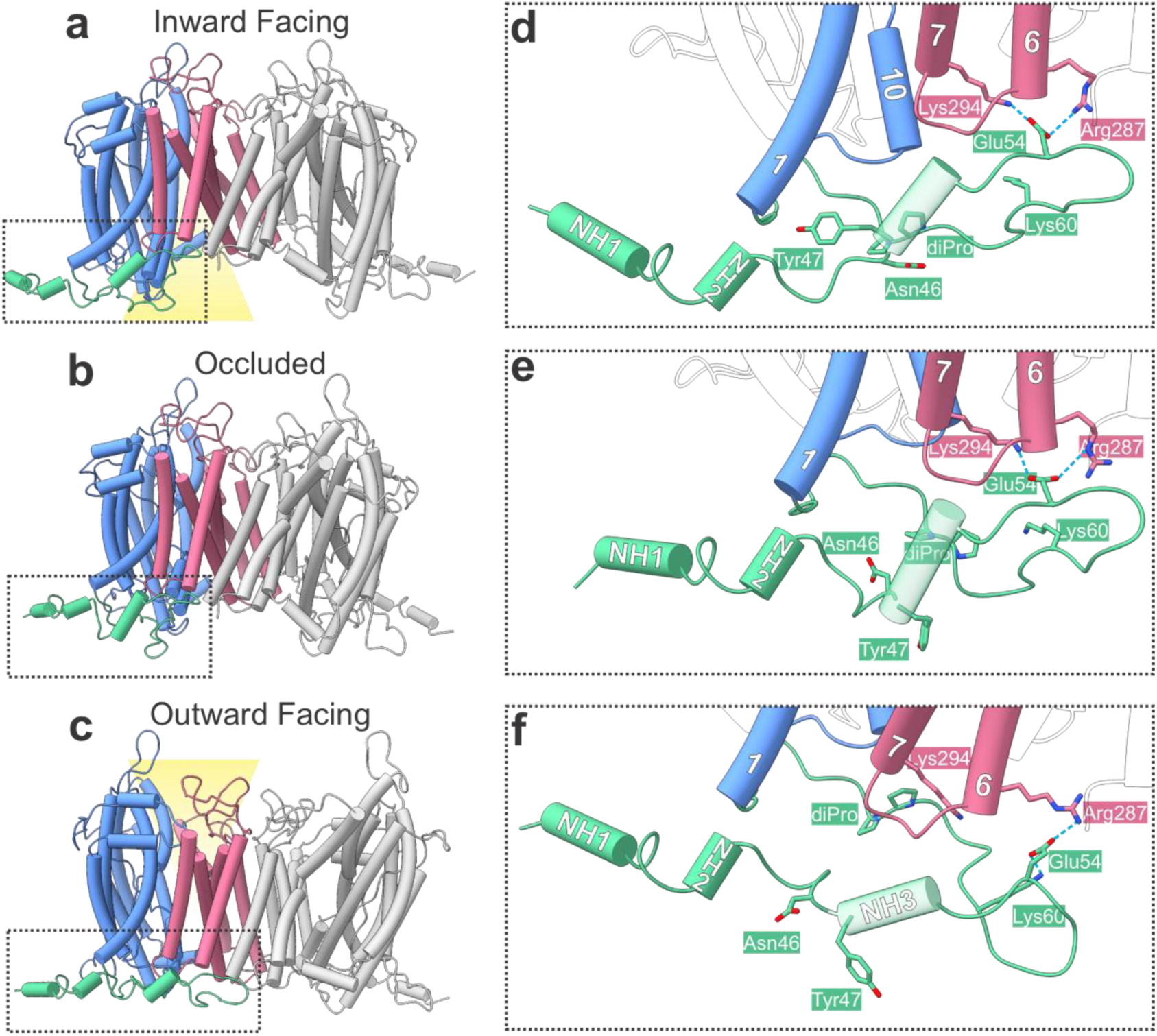
Functionally important N-tail interfaces re-arrange to accommodate sliding of the core domain. **(a-c)** Molecular models of UapA in IF (cryoEM, a), Occ (homology model, b), and OF (homology model, c) states. **(d-f)** Interactions of the N-tail with the core and scaffold domains. Key residues are highlighted.

More specifically, while the stable N-tail-core interactions observed in the IF state (e.g., **Figure 2e**) remain relatively unchanged during the IF-Occ-OF transition, the necessary structural re-arrangements of the N-tail interfaces concentrate on the hydrophobic cavity in the TMS1 of the N-tail-core interface, formed by Leu77, Leu78, and Ile81 (**Figure 2c**), as well as on the bidentate salt-bridge (Arg287-Glu54-Lys294) of the N-tail-scaffold interface (**Figure 2f**). This is particularly important, as our functional studies show that precisely these N-tail structural elements are crucial for controlling UapA transport dynamics. In particular, the IF-Occ transition is characterized by an internal rotation of the core domain, bringing the active site closer to the scaffold (**Figure 3b**). This movement preserves the aforementioned tail interactions but shrinks the hydrophobic pocket of TMS1, affecting Tyr47. Tyr47 disengages, rotates externally, and is replaced by Asn46, another conserved residue occupying the significantly altered pocket (**Figure 3e**). Successively, the Occ-OF transition shows two parallel movements; in the scaffold domain, TMS13-TMS14 approximate the opposing protomer, flattening the interdomain sliding surface, thus allowing the core domain to rotate and extend extracellularly, allowing access to the substrate-binding site (**Figure 3c**). In conjunction, those relative movements increase the distance needed to be covered by the tail, which is stretched, extending the β-sheet-like core interaction facilitated by the local refolding of the diproline motif Pro65-Pro66. Interestingly, the original bidentate salt bridge (Arg287-Glu54-Lys294) is altered, with Lys60 replacing Lys294 and extending the N-tail further (**Figure 3f**). Taken together, those findings suggest a dynamic and complex mechanism of control and modulation of the sliding by the tail, facilitated by numerous interacting surfaces throughout the translocation cycle.

### The high resolution of the substrate binding site rationalizes UapA specificity

The substrate-binding site of UapA is situated halfway through the core domain, sandwiched between the half helices TMS10 cytoplasmically and TMS3 extracellularly. Laterally, it is enclosed by TMS1 and TMS8 (**Figure 1e**, **Figure 4a**). The scaffold, specifically TMS5 and TMS12, acts as a lateral bounding surface and presumably as a sliding surface against which the core domain, and thus the substrate-binding site, moves.

**Figure 4:**
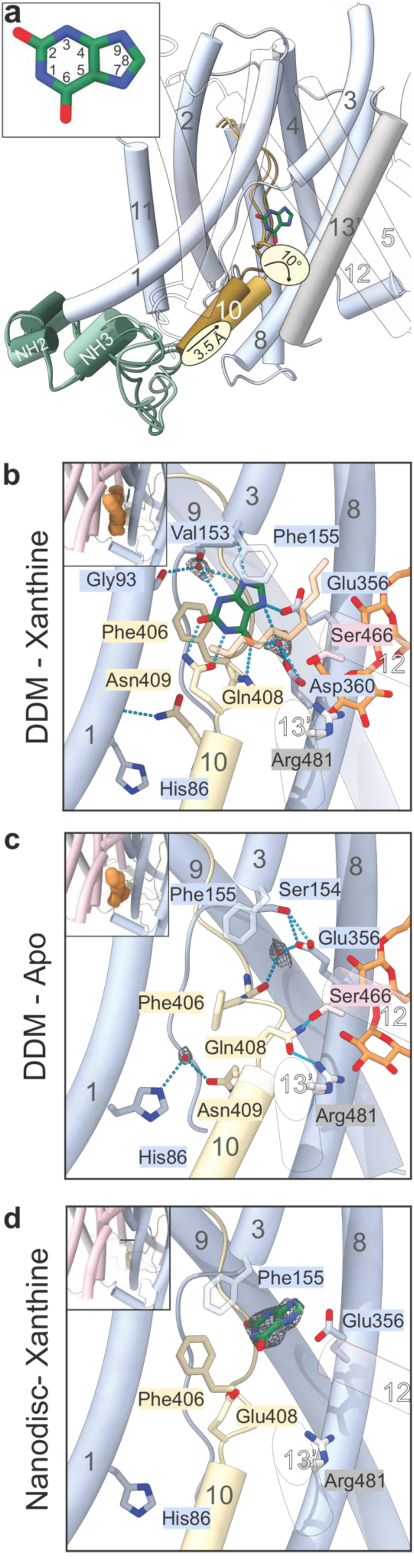
The binding site. **(a)** Overview of the protomer as seen from the side, showing the position of the substrate binding site. The regions with an RMSD > 1Å between the different structures are displayed for each, with the lighter one corresponding to the Apo state and the darker one to the Xanthine-bound state. The core domain is colored light blue. The scaffold domain and the opposing protomer are transparent, except for TMS13 in gray. TMS10 is shown in ochre, and the N-terminal tail is in green. **(b-d)** Closeup of the binding site in the different structures. Side-views can be seen on the top left of each panel, with orange demarcating the lipids detected in the pocket. **(b)** The binding site in UapA_wt_-Xan-DDM. Notice that all polar sites of the xanthine are hydrogen bonded. **(c)** The binding site in UapA_wt_-Apo-DDM. Notice the H-bonds between Gln408 and Arg481 of the opposite protomer and Ser466 of the scaffold domain. **(d)** The binding site in UapA_wt_-Xan-ND. Phe155 is perpendicular and directly above the aromatic Xanthine, securing it in its pocket.

Although the substrate binding site has been previously determined in the crystal structure of mutant G411V at 3.6 Å^4^, we set out to replicate the result in the WT by cryoEM, aiming to improve resolution and provide new detailed insights into substrate recognition. The resulting 2.42 Å UapA_WT_-Xan-DDM reconstruction showed an essentially unchanged configuration of the binding site (**Figure 4b**). However, the significantly improved resolution allowed us to fit several water molecules in the structure, including two directly involved in xanthine binding (**Figure 4b**).

Several intertwined interactions mediate the apparent stability of the substrate; firstly, two prominent Phe residues, Phe155 of TMS3 and Phe406 of TMS10, stack the aromatic ring laterally and proximally, fixing it in place via a triple π-π stacking. Furthermore, all polar sites on the xanthine molecule are hydrogen-bonded directly or via water molecules to the protein. The most stabilizing interaction seems to be that with Gln408, which binds xanthine at C2=O and N1-H in a bidentate manner. A water molecule binds N3-H and N9, with N9 also bound by the peptide nitrogen of the capping Phe155. The water molecule is stabilized by interactions with the backbones of Gly93 (TMS1) and Val153 (TMS3). On the other side, another water molecule is within bonding distance of both C6=O and N7-H, which also interacts with the backbone of Ala407 and the sidechain of Glu356 in TMS8. The water molecule is basically coordinated by the sidechain of Asp360 (TMS8).

Intriguingly, two detergent- or lipid-like densities can be observed near the binding cavity in the xanthine-bound structure. The density in direct proximity to TMS8 can be attributed to a DDM molecule, bound by hydrophobic interactions in the tail and polar ones on the head (**Figure 4b)**. The density directly above the xanthine is indistinct and has been built as a lauric acid, given the absence of head-group information and the resulting clashes involved if built as a DDM (**Figure 4b**).

To assess the role of water molecules and detergent/lipids in substrate binding we first applied a solvent mapping calculation to the xanthine-bound structure. The water molecules surrounding the substrate are predicted a priori, almost in the same position as experimentally defined (**Supplementary Figure 8a**). Omitting them in Induced Fitting Docking calculations (IFD) leads to unfamiliar high-energy results, underlining their necessity for substrate binding. Interestingly, repeating the water mapping analysis upon removing the lipid/detergent molecules revealed high-energy trapped water molecules to be localized precisely in the same positions as the lipophilic detergent side chain. In contrast, stable low-energy waters are located in the positions of the hydroxyl oxygens of the detergent head moiety. This suggests that lipid-like molecules might entropically stabilize the structure in this conformational state, expelling trapped water molecules in the medium and forming H-bonds between hydrophilic heads with the protein (**Supplementary Figure 8b**). Presumably, lipids, probably replaced by DDM molecules during the solubilization process, might lock the binding site in this remarkably stable conformation. We subsequently analyzed xanthine retention in MD simulations. Omitting both water molecules and detergent/lipids results in a cytoplasmic substrate exit after ∼55ns. Adding the water molecules increases that interval to ∼350ns, whereas adding both water and lipid/detergent molecules results in total retention for the total duration of the simulation (400ns). Taken together, these results show an apparent stabilizing effect of both water and lipid/detergent molecules on the active site in the substrate-bound IF structure.

The plethora of interactions described here shows a very stable engagement of the substrate, albeit with varying strengths at each position, rationalizing the high-affinity binding of xanthine by UapA. Given that more polar interactions stabilize the substrate significantly than less polar ones and that water molecules can easily be displaced, we can also rationalize the relatively high, albeit reduced compared to xanthine, affinities to several 3-,7-, and 9- substituted xanthine analogs. For example, 1-methylxanthine, presumably disrupting the core interaction with Gln408, has a *K*_i_ of 280 mM, while 3-methylxanthine, which probably displaces a water molecule, has a *K*_i_ of 28 mM^11^. Similarly, hypoxanthine, guanine, or adenine, all very poorly recognized by UapA (i.e., *K*_i >_ 2mM), lack the C2=O, contrasting the high-affinity binding of UapA for xanthine and uric acid^9^. To calculate the thermodynamics of relevant interactions, we estimated the ΔΔG of hypoxanthine *versus* xanthine binding after IFD using Free Energy Perturbation calculations. The pose was identical to that of the xanthine, but hypoxanthine could not bind Gln408, giving a ΔΔG = +1 kcal⋅mol^-1^, partially explaining the experimentally deduced +3,2 kcal⋅mol^-1^.

We then set out to determine the conformation of the binding site in the absence of xanthine, repeating the same purification and cryo-EM strategy but without including the substrate. This gave rise to the 2.06Å UapA_WT_-Apo-DDM reconstruction with an utterly unobstructed substrate site. Despite this reconstruction also indicating an IF conformation (**Supplementary Figure 9**), we can observe several changes, with the most prominent being the 3.5Å shift and 10° rotation of TMS10 laterally and extracellularly (**Figure 4c**), presumably to fill the void remaining in the absence of substrate. Mechanistically, we consider this structure to represent the conformation after the cytoplasmic release of the substrate, given that there is no extracellular access to the binding site. After a closer inspection of the interactions, the most prominent local change of the pocket in the apo form concerns Gln408 (**Figure 4b, c**). The movement of TMS10 positions it near the differentially charged surface created by Arg481 on TMS13 and Ser466 on TMS12 of the *opposing* protomer. This interaction neutralizes the partial charge caused by the absence of substrate and restabilizes the protein, presumably setting it in a conformation sufficient to be reset to the outward open apo state. Given the absolute conservation of all three residues involved in this interaction in fungal NATs, plus the already established importance of Arg481 for function and specificity^4,44^, we consider this interaction essential for transport activity. Interestingly, Arg481 displays alternative locations for its sidechain, with only one being within hydrogen-bonding distance of Gln408 **(Supplementary Figure 10b)**. However, in the mutant structure UapA_Q408E_-Apo-DDM, Arg481 is clearly positioned only in the hydrogen bond-capable rotamer despite a practically zero overall RMSD, presumably because of the stronger interaction caused by this mutation **(Supplementary Figure 10c)**. This difference can shed light on the cryo-sensitive nature of the mutation. The stronger interaction can increase the energy barrier for a conformational change, thus making it much rarer at lower temperatures.

Glu356, another polar residue in the pocket exposed after xanthine release, rotates extracellularly and interacts with Ser154. In addition, Asn409, previously holding TMS10 into position by interacting with the backbone of His86 in TMS1, is now positioned too far away to meditate on this interaction. Instead, it binds to the sidechain of the same residue through a water molecule (**Figure 4b, c**). As these residues are absolutely conserved and specific relative mutations cause complete loss of transport and/or lead to protein instability, these interactions are crucial to further stabilizing the empty binding site upon substrate release^33,34^. Compared to the xanthine-bound structure, the apo structure shows only the DDM molecule near TMS8 and lacks the lipid-like density above xanthine **(Figure 4b, c).**

We further considered a detergent-free reconstitution system and reconstituted xanthine-loaded UapA in linear nanodiscs supplemented with yeast lipids. Due to processing difficulties regarding the WT in lipid nanodiscs, we opted for the more stabilized mutant, UapA_Q408E_. The resulting reconstruction (UapA_Q408E_-Xan-ND) reached a final resolution of 3.6Å, showing a binding site void of any DDM molecules and lipids and intriguingly, xanthine positioned more extracellularly compared to the WT, less stably wedged between Phe155 and Glu356 (**Figure 4d**). Otherwise, the overall conformation and organization of TMS10 are largely similar to the xanthine-bound conformation in DDM (**Figure 4d**). This indicates that neither lipids nor DDM affects the downward movement of TMS10 in the substrate-bound state. Instead, it is the presence of xanthine that influences this movement. Docking studies on the UapA_WT_-Xan-DDM mutated to Q408E show that the alternate positioning of the substrate is a direct result of the mutation and not of the nanodisc reconstitution and the absence of lipids in the active site. Therefore, we consider this positioning a locked translocation intermediate (**Supplementary Figure 8c**).

### Multiple interactions of reconstituted UapA with lipids/detergents indicate the importance of the lipid environment in transporter functioning

Previous lipidomics experiments on UapA revealed the complex identity of co-purified lipids^6^. Delipidation results in the dissociation of the dimer into monomers, but the subsequent addition of phosphatidylinositol (PI) or Phosphatidylethanolamine (PE) rescues the dimer, indicating that lipid binding is essential for the formation of functional UapA dimers. While previous studies^6,34^ provided evidence that lipid interactions affect both functional dimerization and trafficking, direct evidence on the exact localization of lipids on the protein, their functional importance at those sites, and their impact on the successful assembly of functional UapA dimer remained unclear. The high-resolution cryoEM maps of UapA_WT_-Apo-DDM, UapA_WT_-Xan-DDM, and UapA_Q408E_-Apo-DDM provide the first detailed structural insights into UapA-lipid interactions, revealing at least 80 elongated densities that can be attributed either to DDM detergent moieties or lipids that co-eluted during detergent-mediated purification (**Figure 5a**). Most of these decorate the hydrophobic surface of the protein peripherally, forming an annulus around the UapA dimer, and are crucial for its stabilization in the membrane and, apparently, for trafficking *in vivo*. Importantly, we also observe high-affinity interactions deeply buried within the protein structures, either in the binding site or the dimerization interface. These interactions are of particular interest, as they may play critical roles in UapA structure and function, as our previous genetic and biochemical analysis supports. Notably, the three high-resolution cryoEM structures of UapA extracted by DDM are almost indistinguishable regarding densities attributed to lipid/detergent molecules. For clarity, **Figure 5a** illustrates the distribution of lipid/detergent molecules in UapA_WT_-Xan-DDM, but the following description applies to all three cryoEM structures.

**Figure 5:**
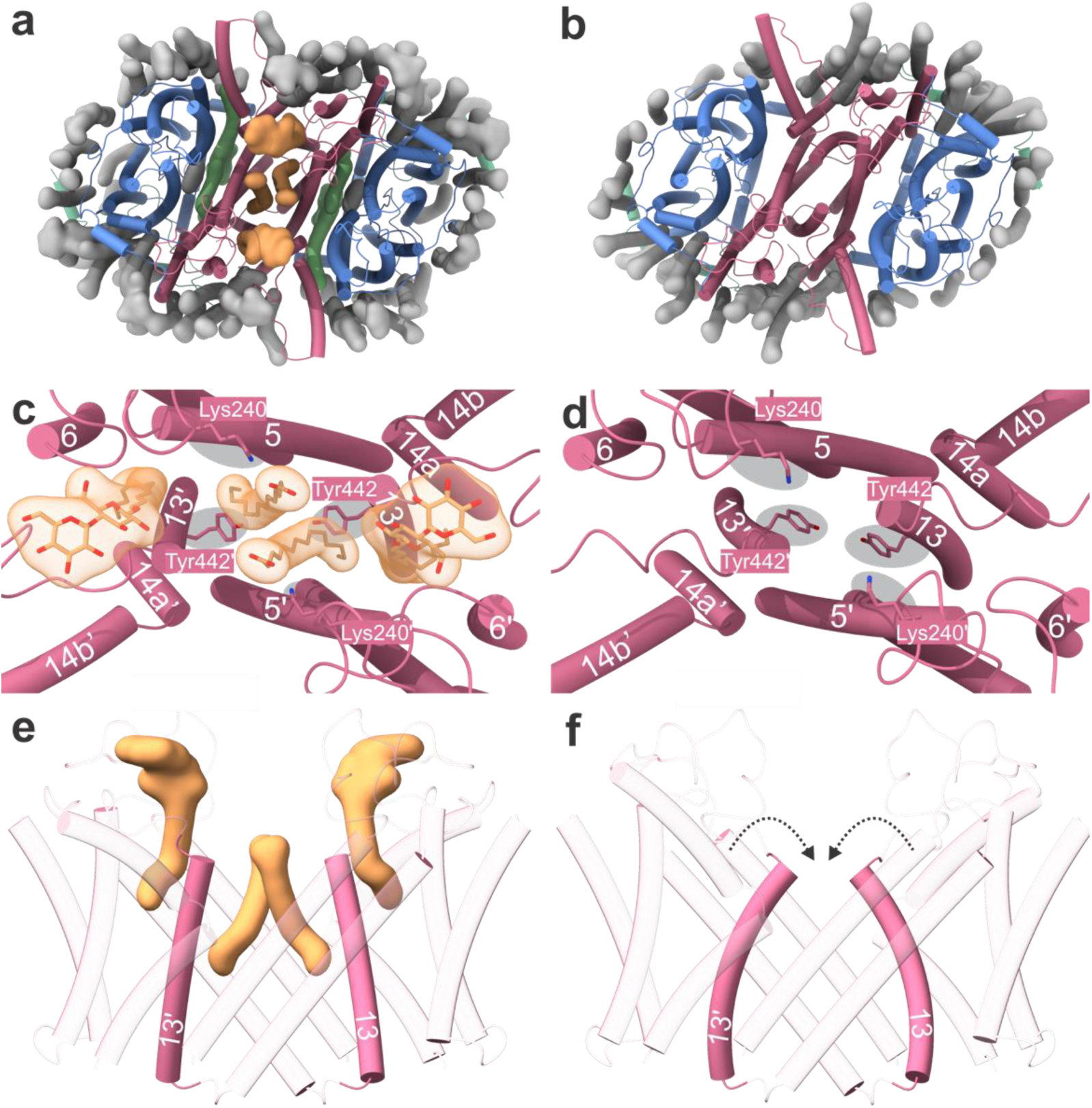
Lipid localization. (**a-b**) Location of densities attributed to different lipid or detergent species in UapA_WT_-Xan-DDM (a, DDM) and UapA_Q408E_-Xan-ND (b, nanodisc). Annular lipids are colored gray, binding site lipids are colored green, and dimerization lipids are colored orange. **(c-d)** Dimerization interface in DDM (c) structure and nanodisc structure (d) **(e-g)** Side view of the Scaffold domains in the dimerization interface in DDM (e) and nanodisc structure (f). Notice the movement of the extracellular end of TMS13.

Elucidating the exact identity of these species was challenging in most cases despite high average resolution. We could unequivocally assign densities as lipid or detergent at several positions by observing two connected fatty acid chains or finding chain lengths inconsistent with DDM molecules (**Supplementary Figure 11a,b**). However, determining the exact identity of the lipids was often not feasible due to significantly lower local resolution in the head-group region. Therefore, we modeled ambiguous densities as myristic acid.

The annular lipids surrounding the transporter are mostly non-descript, with little or no headgroup information (**Figure 5a**; gray). A notable exception is a DDM molecule bound stably near the previously identified tentative lipid-binding amino acids Lys73, Arg133, and Arg421 on the contralateral edges of the protein (**Supplementary Figure 11c**). In addition to several fatty-acid-like densities, we observed several ergosterol molecules present in fungal membranes instead of their mammalian equivalent, cholesterol. The most notable example is an ergosterol molecule stably bound by Trp330 on top of TMS9 of the core domain (**Supplementary Figure 11d**). This position is conserved as Trp or Phe in eukaryotic NATs but not in prokaryotic, which lack sterol in their membranes. The intriguing lipid densities identified in the binding site (**Figure 4b-c and Figure 5a**; green) and their effect on substrate binding are described in detail in the previous section (see **Figure 4**).

Importantly, we identified two pairs of lipid/detergent molecules (two molecules per monomer) directly positioned at the interface between the two monomers on the cytoplasmic face of the transporter. The first one binds tightly at the interface formed between TMS12 and TMS6 of the scaffold domain of one protomer to TMS13 of the scaffold domain of the opposing monomer (**Figure 5c**, lateral detergent molecules). These densities were attributed to DDM molecules because of the length of the acyl chain and the head group bulkiness. Surprisingly, we also observe two phospholipids at the exact center of the dimerization interface between the scaffold domains of the opposing monomers, buried deeper than all other lipids in the structure and completely protected from the bulk membrane lipids **(Figure 5c**, central lipids). In addition to hydrophobic interactions along their tails, they are held in place by two residues on each protomer, Tyr492 and Lys240. To test the importance of these residues, we analyzed respective Ala substitutions. While Ala replacement of Tyr492 had no distinct effect on trafficking and transport activity of the UapA, Ala replacement of Lys240 led to complete trafficking arrest at the ER, protein instability (e.g., sorting to vacuoles) and undetectable function (no growth on uric acid or xanthine) (**Supplementary Figure 7**).

To further investigate UapA-lipid interactions in a more native lipid environment, we analyzed the cryoEM structure of UapA_Q408E_-Xan-ND reconstituted in lipid nanodiscs supplemented with yeast polar lipid extract (**Figure 5b**). This structure was expected to replicate protein-lipid interactions similar to those found *in vivo*^45^, although the protein was initially solubilized by detergent. The distribution of annular lipids around the protein in the nanodisc structure closely resembles that in the three structures obtained in detergent (**Figure 5a-b**, gray). Those interactions serve thus as stable anchor points for the protein in the membrane. Surprisingly and in stark contrast to the three detergent structures, the UapA nanodisc structure did not include any buried lipids at either the dimerization interface or the binding site (compare **Figure 5a** to **5b**). The absence of these lipids resulted in a distinct dimer conformation. Specifically, the lack of central lipids causes the cytoplasmic side of TMS13 to tilt and approximate its protomeric counterpart. This movement brings the Tyr492 residues of each TMS13 closer, closing the gap left by the missing interactions with the central lipids (**Figure 5d, f**). This re-arrangement leads to a slight clockwise rotation of the two protomers by approximately 10°, resulting in a more compact dimer conformation with an overall width reduction of 2.2 Å. The reasons for the absence of specific buried lipids in the nanodisc structure and the mechanism behind this partial yet significant delipidation remain unclear. It is important to note that achieving the nanodisc structure required extensive processing and eliminating a large portion of the dataset, resulting in a significantly lower resolution in the final reconstruction than the detergent structures. Furthermore, the two monomers in the nanodisc structure are not uniformly resolved, reflecting a higher degree of flexibility likely due to the weaker dimeric interface. A leading hypothesis is that the biobeads might have absorbed the buried lipids during the nanodisc assembly process, leading to a more compact protein conformation of the dimer. However, despite the overall compactness, the dimeric interface notably weakens. Nevertheless, UapA maintains its dimeric assembly in the membrane environment, even without these central buried lipids.

## Discussion

Under the light of the present highly resolved structures, coupled with a long history of genetic and functional analysis of UapA, we unveiled several unprecedented aspects of the elevator-type transport mechanism of a eukaryotic NAT transporter. Of particular importance, we managed to address two significant issues for the first time. First, how an extended cytosolic N-tail region, which is absent in several prokaryotic NAT homologues or highly variable in other eukaryotic NAT members, might affect UapA function. Second, how UapA high substrate specificity is determined. Our work shows that the N-tail forms a tightly bound lid interacting with both the core and scaffold domain via multiple polar and hydrophobic bonds. Previous and new mutational analysis has confirmed the importance of these interactions in both proper ER exit and transport activity. MD has further suggested that the N-tail remains well-bound to the core and scaffold domains during the transport cycle. Supported by our extensive genetic analysis, such a tight interaction of the N-tail with the transporter core and scaffold domains seems to stabilize a conformation necessary for ER exit and trafficking to the PM. UapA, as all transmembrane proteins, needs to be recognized by the COPII machinery and packaged into nascent vesicles (or extended tubules) budding from specific ER-Exit Sites (ERES), which eventually form carriers for translocation to the PM^46,47^. Although the current consensus is that secretory or membrane proteins targeted to the PM possess short linear ER-exit motifs, this is by far an oversimplification as no such motifs exist on most polytopic transmembrane proteins and, in particular, solute transporters. In addition, the few motifs described are context-dependent, as their transfer to another protein cannot promote ER exit^17^. Thus, the mechanism of ER exit of transmembrane cargoes, such as UapA and other transporters, remains elusive. This study and previous genetic analysis on UapA strongly suggest that the ‘code’ for proper, COPII-dependent ER-exit is the fine 3D folding of cytoplasmic-facing transporter domains, with a prominent role of the N-tail domain. In line with this concept, unbiased genetic approaches to select specifically ER-retained mutants of UapA led to mutations that are located very often on the cytoplasmic side of TMSs or internal loops (E. Demos, S. Dimou and G. Diallinas, unpublished). Overall, the presently described structure of the N-tail and its associated interactions with cytosolic-facing ends of TMS or other internal loops strongly suggest that the ‘code’ needed for recruitment of UapA to ERES, packaging into COPII carriers and ER-exit is the proper 3D folding of cytoplasm-exposed domains of the transporter. As the cytosolic domains of polytopic transmembrane proteins interact, in principle, with lipids of the inner side of the membrane bilayer, specific lipids should also play an essential role in correct folding and proper ER exit of the transporter. Along these lines, our present study revealed multiple interactions of the UapA N-tail and other internal segments with detergents and lipids. In conclusion, we believe that neosynthesized UapA, rapidly after its co-translocation into the ER, dimerizes and acquires a specific and relatively compact conformation through interactions of its N-tail and other cytosolic domains with ER lipids. This compact structure seems necessary for partitioning into ERES, where UapA interacts with COPII components to be appropriately packaged into vesicular trafficking carriers. Upon integration of UapA to the PM, which has a very different lipid composition and width compared to the ER membrane, interactions with lipids may be modified so that the compact UapA structure might be loosened to account for more flexible, transport-competent, alteration of conformations, necessary for transport catalysis (see also later).

Eukaryotic transporters, in general, are characterized by long cytosolic tails, most often absent in their prokaryotic homologs. Animal and plant homologs of UapA also have long cytosolic tails, similar to all fungal NATs. Importantly, in several cases, transporter tails have been shown to affect not only trafficking, cellular expression, and turnover, as might have been expected, but also transport function and, surprisingly, specificity^30,48,49^. Somewhat paradoxically, bacterial homologs of UapA that practically lack extended cytosolic N-tails (e.g., UraA) show very similar transport kinetics and specificity to UapA. In the simplest scenario, the tails of transporters evolved in eukaryotes under the challenge of subcellular trafficking and the need for posttranslational cellular regulation. However, the evolution of longer tails should necessarily be adapted or even integrated into the transport mechanism. This, in turn, allowed tails to acquire new roles in transport catalysis. The ideas above align with the observation that the tails of homologous transporters are by far more variable than their transmembrane segments, indicating a very different rate of evolution.

Concerning UapA specificity, the high resolution of the substrate binding site, including mapping of water molecules, explained why UapA is a high-affinity transporter for uric acid and xanthine but transports other purines very poorly. For example, UapA has a *K*_m_ for xanthine close to 7 μΜ, while the affinity for hypoxanthine or other purines is >300-fold lower (*K*_i_ > 2 mM). We showed that xanthine binds to UapA with interactions of all its polar sites, directly or via three water molecules, and additionally is stacked by two Phe residues. Hypoxanthine and other purines lack C2=O and have distinct protonation states in N3, N7, or N9. Thus, they are less stabilized in the substrate binding site. Similar observations can now be extended to purine and pyrimidine analogs, including first-line antifungals or other purine-based drugs that bind with differential affinities to UapA.

Our studies have yet to identify an OF conformation of UapA. Intriguingly, this is a common theme in all dimeric elevator transporters resembling the structure of NATs. Most reported structures, except the chloride/bicarbonate anion exchanger Band3 and the CNT3 purine transporter, are either IF or partially occluded. The OF conformations of Band3 and CNT3 have been obtained by ‘locking’ the transporters with a conformation-‘trapping’ inhibitor^29,50^. Purified NAT-like elevator transporters seem to be hardly stable in the OF conformation. Given the uniqueness of the genetic analysis of transporters in *A. nidulans*, the long-sought outward conformation of NAT proteins, necessary to establish a complete transport cycle mechanism, might be achieved using specific UapA mutations that would lock the transporter in this relatively unstable conformation.

Interestingly, a recently published cryoEM study on hSVCT1 reports the IF state, including the interactions with lipids/detergents and a unique new conformation, a substrate-free Occ state, obtained solely when the transporter was purified in the absence of the naturally symported Na^+^ ions^22^. In the IF state of hSVCT1, lipids/detergents are mainly positioned on the surface over the inter-protomer interface, whereas in our IF UapA DDM structure, lipids/detergents are in addition, directly positioned and buried at the exact center of the interface, stabilizing the dimer. Thus, while lipid-protein interactions significantly contribute to oligomerization and transporter activity, different NATs exhibit distinct protein-lipid interactions at their dimer interface. This feature might be essential for fine-tuning their transport properties. Regarding the unprecedented substrate-free occluded state of hSVCT1, the authors of this work consider that this new conformation represents a structure formed while the substrate-free IF returns to OF states. In this state, the entire scaffold domain tilts toward the cytoplasmic side of the core domain in the same protomer, with TMS6 and TMS13 rotating outwardly so that the scaffold/dimerization domain undergoes a rigid rocking bundle-like movement, similar to the one prosed in other transporters, but also speculated for UraA, a bacterial NAT^20^. A consequence of this movement of the scaffold domain is a dramatic modification of the dimerization interface, which loosens and becomes invaded with lipids/detergents. The authors speculate that *in vivo,* this invasion of lipids elicits the formation of the OF conformation as a natural step in the transport cycle.

In our nanodisc IF UapA structure, delipidation causes significant compression and simultaneous weakening of the dimeric interface, involving a major rotational shift of the complete monomers. Despite the compression, the two scaffold domains exhibit looser associations without central lipids, potentially allowing enhanced independent monomer action and/or lipid-dependent rearrangements that may facilitate drastic domain motions in the transport cycle, similar to SVCT. In contrast, lipid binding at the dimer center stabilizes the scaffold interface and might promote cooperativity and coordinated action between the monomers during transport. Whereas such speculations have to be experimentally confirmed, overall, the lipid-dependent structural flexibility of the central interface in UapA and the recent insights from the substrate-free Occ state of SVCT1 suggest that transport activity in the NAT family of transporters might not only depend on lipid-mediated dimerization but also on fine lipid-dependent rearrangements of the dimeric interface which thus determines the plasticity of transport dynamics and substrate specificity. Further investigation is needed to determine whether such dynamic re-arrangements of central lipids in the dimer interface fine-tune monomer cooperativity, transporter activity, and specificity.

## Acknowledgments

Work in the laboratory of G.D. was financed by an HFRI grant (KE 18458). Yiannis Pyrris was supported by a Foundation Santè fellowship. The cryo-EM data were collected at “Cryo-EM SoN”, the cryo-EM infrastructure of the University of Münster, funded by the Deutsche Forschungsgemeinschaft (DFG, German Research Foundation) – Projectnumber 496113311. The cryo-EM dataset was processed at the Palma II HPC (DFG INST 211/667-1) of the University of Münster. We acknowledge technical support from the HPC team at the University of Münster. The authors kindly thank Prof. Bernadette Byrne (Imperial College, London) for providing the UapA_wt_ and UapA_Q408E_ plasmids and expressions strain, and for general guidance with the purification procedure. Molecular graphics and analyses performed with UCSF ChimeraX, developed by the Resource for Biocomputing, Visualization, and Informatics at the University of California, San Francisco, with support from National Institutes of Health R01-GM129325 and the Office of Cyber Infrastructure and Computational Biology, National Institute of Allergy and Infectious Diseases.

## Materials and Methods

### Purification of UapA

UapA_WT_ and UapA_Q408E_ UapA were purified from *Saccharomyces cerevisiae FGY217 (MATa, ura3-52, lys2-201, and pep4),* harboring a pDDGFP-2 plasmid containing UapA cDNA, under a galactose-inducible GAL1 promoter, and the purification was performed largely as described previously^4,40,51^. All cultures were done at 30°, 300 rpm (Infors HT Multitron). A single colony was used to inoculate a 10ml SC-Ura (2% glucose) culture overnight (16h). The next day, the preculture was used to inoculate a 400ml SC-Ura (2% glucose, 6.7 g/L yeast nitrogen base without amino acids, 2 g/L amino acid mix without uracil) preculture overnight (16h). The final preculture was used to inoculate 6-12L pre-warmed SC-Ura (0.1% glucose) at an initial OD_600_=0.12 in baffled flasks and incubated until OD_600_=0.6 (∼6h). Galactose was added to a final concentration of 2% from a 25% galactose, 1x SC-Ura stock, and the cultures were allowed to incubate for a further 24 hours. Cells were harvested by centrifugation (4,000g, 10 min) (JLA 8.1000, Beckmann Coulter). All the following steps were performed in ice in a 4°C cold room, with pre-chilled buffers and rotors. Cells were resuspended by pipetting in 10 mL cell resuspension buffer (50 mM Tris, 1 mM EDTA, 600 mM sorbitol, 1 cOmplete protease inhibitor tablet EDTA-free per 50ml, pH=7.5) per L cell culture. Lysis was induced by passing the resuspended cells through a Microfluidics LM10 Microfluidizer 10 times under 23.000 psi while keeping the temperature below 10 °C. The lysate was further centrifuged to pellet unbroken cells and cell fragments (10,000g, 10 min, 4°C) (JA-25.50, Beckmann Coulter). The supernatant was ultracentrifuged to collect membranes (150,000g, 2h, 4°C) (Type 45 Ti, Beckmann Coulter). The pellet was resuspended in 5ml membrane resuspension buffer (20 mM Tris, 0.3 M sucrose, 0.1 mM calcium chloride, cOmplete protease inhibitors, pH=7.5) per L cell culture, and homogenized using a 40ml douncer.

Membrane resuspensions were solubilized by diluting in 100ml/6L solubilization buffer (final concentration after dilution: 1x PBS, 100 mM NaCl, 10% glycerol, 1% n-dodecyl-b-D-maltoside (DDM), cOmplete protease inhibitors, pH=7.5) and stirring for 1hr. The remaining insoluble material was pelleted by ultracentrifugation (100,000g, 45 min, 4°C) (Type 45 Ti, Beckmann Coulter).

The solubilized membranes were incubated for 2 hours with 1ml/L Ni-NTA agarose beads (PureCube, Cube Biotech) pre-equilibrated with affinity buffer (1x PBS, 100 mM NaCl, 10 mM imidazole, 10 % glycerol, 0.03 % DDM, pH=7.5) under very slow stirring. The slurry was then loaded onto a glass Econo-column chromatography column (Bio-Rad) and was washed with 10 column volumes (CVs) affinity buffer followed by 30 CVs affinity buffer supplemented with 30 mM imidazole. UapA was recovered from the resin with 5 CVs affinity buffer, supplemented with 250 mM imidazole.

The elution was used directly for nanodisc assembly. For CryoEM samples, after elution, the protein was concentrated to 250μl using a 100kDa molecular weight cut-off filter (Amicon), centrifuged to pellet aggregates (10,000g, 10min, 4°C) (5418R, Eppendorf) and injected in a Superdex 200 10/300 gel filtration column (Cytiva) pre-equilibrated in SEC Buffer (20mM Tris, 150 mM NaCl, 0.03% DDM, pH=7.5) on an NGC Medium-Pressure Chromatography System (Biorad). The highest concentration fractions were concentrated to 4-6 mg/ml, quantified using the Micro BCA Protein Assay Kit (Thermo Scientific) and plunged directly. Neither the cells nor the protein was frozen at any point during purification. For preparations containing xanthine, it was added to all purification buffers and media at a concentration of 600 μM.

### Purification of MSP1E3D1

pMSP1E3D1 was a gift from Stephen Sligar (Addgene plasmid #20066; http://n2t.net/addgene:20066; RRID: Addgene 20066). Protein expression and purification were done as described previously^52–54^. In short, the plasmid was transformed into competent *E. coli* BL21(DE3) cells and plated on an LB-kanamycin plate. Transformants were inoculated in 30 ml LB supplemented with kanamycin overnight (37°C, 220 rpm). The overnight culture was added to 1 L Terrific broth supplemented with kanamycin and incubated (37°C, 220 rpm) until an OD_600_ of 2.5 was reached. Protein expression was induced by supplementing the medium with 1 mM IPTG. After 4h, the cells were harvested (4000g, 30min, 4°C) (JA-25.50, Beckmann Coulter) and resuspended in 25ml cell resuspension buffer (20 mM sodium phosphate (pH 8), 1 % Triton X-100, cOmplete tablet, 10μg/ml DNase I, 0.2mg/ml lysozyme). Cells were stirred for 30min at 4°C, and then passed once from a Microfluidics LM10 Microfluidizer, under 23.000 psi. The crude lysate was then centrifuged to remove inbroken cells and cell debris (40,000g, 30 min, 4°C) (JA-25.50, Beckmann Coulter). The supernatant was loaded on an Econo-column (Biorad) containing 2.5ml Ni-NTA agarose beads (PureCube, Cube Biotech), pre-equilibrated with cell resuspension buffer. The lysate was allowed to pass through, and the material was washed with 10 CV wash buffer A (40 mM Tris (pH 8), 300 mM NaCl, 1 % Triton X-100), 10 CV wash buffer B (40 mM Tris (pH 8), 300 mM NaCl, 20 mM imidazole, 50 mM sodium cholate), 10 CV wash buffer C (40 mM Tris (pH 8), 300 mM NaCl, 20 mM imidazole), and finally 10 CV wash buffer D (40 mM Tris (pH 8), 300 mM NaCl, 50 mM imidazole). MSP was eluted using 5 CV elution buffer (40 mM Tris (pH 8), 300 mM NaCl, 300 mM imidazole) and loaded onto a desalting column equilibrated with PD-10 buffer (50 mM Tris (pH 7.5), 100 mM NaCl, 0.5 mM EDTA). 1 OD_280_ unit of in-house His-Tagged TEV protease was mixed with 100 OD_280_ units of MSP and allowed to react overnight at 4°C. The mixture was then passed again through the Econo-column, pre-equilibrated with cell resuspension buffer, and washed for 5 CV with buffer D. The flowthrough was collected, and the tagless MSP was concentrated to 10 mg/ml using a 30kDa MWCO filter (Millipore), flash-frozen, and stored at −80 °C.

### Reconstitution of UapA in Nanodiscs

S. cerevisiae polar extract lipids (Avanti Polar Lipids) were dissolved in chloroform at 25 mg/ml. A lipid film was generated by evaporating the chloroform under vacuum. Liposomes were suspended in lipid buffer (20 mM Tris (pH 7.4), 150 mM NaCl, 7.5 % DDM (w/v)) at a concentration of 40 mg/ml followed by sonication in a water bath for 30 minutes and three cycles of freeze-thawing using liquid nitrogen. Lipids were stored at −80 °C or used directly. The nanodisc mixture, consisting of UapA, MSP1E3D1, and lipids in a molar ratio of UapA:MSP: lipids = 1:10:1250, was incubated overnight at 4°C on a rotating mixer. Biobeads SM-2 (Bio-Rad) were hydrated by washing them once in methanol, followed by three washes in distilled water. 1 mg Biobeads was used for every μg of UapA in the nanodisc mixture to remove the DDM. ¼ of the total quantity of Biobeads was added to the mixture every hour for a total of 4 hours. The final addition was done at room temperature. Biobeads were pelleted by centrifugation at 1000xg for 1 minute, and the supernatant was aspirated. The Biobeads were washed twice with Nanodisc Buffer (20 mM Tris, 150 mM NaCl, 0.6 mM xanthine, pH=7.5). The sample and washings were injected in a 1 ml HisTrap HP column (Cytiva) and washed with 30 ml Nanodisc Buffer supplemented with 15mM imidazole, to wash away empty nanodiscs. Finally, the UapA-containing nanodiscs were eluted in 10 ml Nanodisc Buffer supplemented with 500mM imidazole and concentrated to 250 μl. The sample was run on a Superdex 200 10/300 gel filtration column pre-equilibrated with Nanodisc Buffer. SDS-PAGE identified fractions containing nanodiscs with UapA embedded and then concentrated to 6-8 mg/ml UapA using a 100kDa molecular weight cut-off filter (Amicon).

### Grid Preparation and Data Collection

For cryo-EM sample preparation, 4 μl protein solution was applied to a glow-discharged UltrAuFoil R 1.2/1.3, 300 mesh (Quantifoil Micro Tools). The grids were blotted for 4.5s and plunge-frozen in liquid ethane after 1 second of drain time using a blot force of −1, in 100% relative humidity at 13°C (Vitrobot Mark IV, Thermo Fischer Scientific). The plunged EM grids were clipped and used for automated dataset collection using the EPU software with a Krios G4 Cryo-TEM (Thermo Fischer Scientific) at 300 kV, equipped with an E-CFEG, a Selectris X Energy Filter at a slit width of 10eV, and Falcon 4i Direct Electron Detector.

### CryoEM Data Processing

Movies for all datasets were monitored for quality and preprocessed on CryoSPARC Live. The gain-corrected, motion-corrected, dose-weighted, and CTF-estimated micrographs were exported for further processing to CryoSPARC (v3.1.0-v4.5.1). For all subsequent jobs described, the following modifications were made to the default settings:

#### 2D Classification

class2D_max_res and class2D_max_res_align were usually increased to 9 and 12 respectively, although decreased to 2 for the final job of the particle stack, class2D_force_max was set to false, class2D_num_full_iter was set to 2-4, class2D_num_full_iter_batch was set to 40-100, and class2D_num_full_iter_batchsize_per_class was set to 400-1000.

#### Ab-Initio Reconstruction

abinit_max_res was decreased to 6-9, abinit_init_res was reduced to 15-25, abinit_minisize_init was set to 300, abinit_minisize was set to 1000.

#### Non-Uniform Refinement

refine_res_init was usually set to 6-12, and refine_defocus_refine and refine_ctf_global_refine were set to true in all datasets but the nanodisc one. refine_do_ews_correct was set to true, given prior knowledge of our microscope.

For the UapA_WT_-Apo-DDM Dataset, 7,5 million particles were blob picked and extracted in a roughly 400Å box at a pixel size of 2.58Å/pix. The particles were subjected to two rounds of strict 2d Classification. The remaining 1 million particles were re-centered and re-extracted at a pixel size of 1,05Å/pix and used for a 3-class Ab-Initio reconstruction, followed by a heterogenous refinement. The largest class showed very high-resolution features at this stage, including side-chain information. The particles belonging to this class were recentered and reextracted at a pixel size of 0,76Å/pix, non-uniformally refined under C2 symmetry. The following 3D alignments were further refined by C2 symmetry expansion and local refinement with a step of 0.05Å, giving the 2,06Å reconstruction (Supplementary Figure 1).

For the UapA_WT_-Xan-DDM Dataset, the procedure was equivalent to the one described above, but templates created from the previous high-resolution map were used for template picking (Supplementary Figure 2).

The UapA_Q408E_-Apo-DDM Dataset and the UapA_Q408E_-Xan-ND Dataset were more challenging, and we used seed-facilitated 3D classification, as described by Wang et al.^21^ to rescue as many particles as possible to get high-resolution structures (Supplementary Figure 3,4).

All models were subjected to model-free density modification and/or anisotropic sharpening using local_aniso_sharpen and resolve_cryo_em from Phenix^55^, respectively.

### Model Building

Models were manually built in ChimeraX v1.8^56–58^ using the Isolde plugin^59^. The initial model for all the structures was the Alphafold prediction AF-Q07307-F1^60,61^. Water molecules were built using phenix.douse from Phenix^55^, followed by manual inspection, including deletion, addition, and relaxation of water molecules in Isolde. Models were real-space refined and assigned b-factors using phenix.real_space_refine.

### Media, strains, and growth conditions for Aspergillus nidulans

Standard complete (CM) and minimal media (MM) for *A. nidulans* growth were used. Media and supplemented auxotrophies were used at the concentrations in http://www.fgsc.net. Glucose 1 % (w/v) was used as a carbon source. 10 mM sodium nitrate (NO_3_) or 10 mM ammonium tartrate were standard nitrogen sources. Uric acid or xanthine was used at the following final concentrations at 0.5 mM. All media and chemical reagents were obtained from Sigma-Aldrich (Life Science Chemilab SA, Hellas) or AppliChem (Bioline Scientific SA, Hellas). A *ΔfurD::riboB ΔfurA::riboB ΔfcyB::argB ΔazgA ΔuapA ΔuapC::AfpyrG ΔcntA::riboB pabaA1 pantoB100* mutant strain, named Δ7, was the recipient strain in transformations with plasmids carrying UapA alleles based on complementation of the pantothenic acid auxotrophy *pantoB100*. A *pabaA1* (paraminobenzoic acid auxotrophy) is a wild-type control strain. A. nidulans protoplast isolation and transformation were performed as previously described^62^. Growth tests were performed at 25 or 37 ° C for 48 h, at pH 6.8.

### Standard molecular biology manipulations and plasmid construction

Genomic DNA extraction from *A. nidulans* was performed as described in FGSC (http://www.fgsc.net). Plasmids, prepared in *E. coli*, and DNA restriction or PCR fragments were purified from agarose 1% gels with the Nucleospin Plasmid Kit or Nucleospin ExtractII kit, according to the manufacturer’s instructions (Macherey-Nagel). Standard PCR reactions were performed using KAPATaq DNA polymerase (Kapa Biosystems). PCR products used for cloning, sequencing, and re-introduction by transformation in *A. nidulans* were amplified by a high-fidelity KAPA HiFi HotStart Ready Mix (Kapa Biosystems) polymerase. DNA sequences were determined by VBC-Genomics. Site-directed mutagenesis was performed according to the instructions accompanying the Quik-Change® Site-Directed Mutagenesis Kit (Agilent Technologies, Stratagene). The principal vector used for most *A. nidulans* mutants is a modified pGEM-T-easy vector carrying a version of the *gpdA* promoter, the *trpC* 3’ termination region, and the *panB* selection marker. Mutations were constructed by oligonucleotide-directed mutagenesis or appropriate forward and reverse primers (see Supplementary Table 3).

### Epifluorescence microscopy

Samples for standard epifluorescence microscopy were prepared as previously described^46^. In brief, sterile 35 mm l-dishes with a glass bottom (Ibidi) containing liquid minimal media supplemented with NaNO_3_ and 1% glucose were inoculated from a spore solution and incubated for 16 h at 25°C. The images were obtained using an inverted Zeiss Axio Observer Z1 equipped with an Axio Cam HR R3 camera. Image processing and contrast adjustments were made using the ZEN 2012 software. Further processing of the TIFF files was done using Adobe Photoshop CS3 software for brightness adjustment, rotation, alignment, and annotation.

#### Homology modelling

The inward cryoEM structure combined with Band3 (PDB ID: 4YZF)^20^ and UraA (PDB ID: 5XLS)^63^ crystallographic structures were used for the development of an Outward and an Occluded model of UapA respectively. The models were obtained utilizing Protein Structure Alignment^64^ and Crosslink Proteins tools^65^ as embedded in Schrodinger Maestro according to the following process. The experimentally defined CryoEM IF monomer structure was divided into two segments, namely the core containing TMS1, 2, 3, 4, 8, 9, 10, 11 and the scaffold containing TMS5, 6, 7, 12, 13, 14 helices of the symporter. This approach was selected to preserve the experimental secondary structure of UapA to the greatest possible extent. The OF model was created by cutting the CryoEM structure into 8 different locations. Specifically, the bonds between Tyr45-Asp46, Pro65-Pro66, Val209-Pro210, Pro220-Ile221, Gly313-Tyr314, Lys332-Thr333, Ile436-Phe437 and Pro448-Asn449 were disrupted. Subsequently, the core helices and the N-terminal parts Gln26-Tyr45 and Pro66-Val76 were aligned with the core domain of 5XLC, while the scaffold domain was superimposed onto the scaffold domain of 5XLC using the Protein Structure Alignment subroutine. The hinge segments Pro210-Pro220, Tyr314-Lys332, and Phe437-Pro448, along with the N-terminal Asp46-Pro65 loop, were manually positioned to facilitate bond formation between the core and gate domains. The secondary structure of the hinges, especially the helices and the Leu50-Pro62 loop, were preserved in the best possible way. Bond formation was achieved by deleting 1 to 5 residues around the interrupted bonds, and missing loop creation was accomplished utilizing the Crosslink Proteins tool.

Occ structure model creation followed the same workflow with minor differences. The structure was split into 2 segments by breaking the bonds between Ile42-Gly43, Leu50-Phe51, Lys60-Asp61, Pro66-Phe67, Phe208-Val209, Ala323-Pro324, and Ile436-Phe437. The core helices and the N-terminal parts Gln26-Ile42 and Pro66-Val76 were aligned with the core helices of 4YZF, while the scaffold domain and the Leu52-Pro62 loop were superimposed onto the scaffold domain of 4YZF. The hinges were positioned without requiring relocation, enabling bond formation between the domains. Missing loops and bond formation were accomplished using the Crosslink Proteins tool, following the same method described earlier. Protein structure alignment was conducted using protein backbone atoms, and the Crosslink Proteins tool was employed with default parameters for all calculations.

#### Docking Calculations

##### Glide Docking

The CryoEM structures of UapA, as well as the OF and Occ models, were processed and minimized using the Protein Preparation Wizard tool^66^ within Maestro (Schrodinger Inc.). Xanthine was prepared using LigPrep^67^ (Schrodinger Inc.), and multiple docking grids were created with multiple combinations of Van der Waals (VdW) radii 0.8 and 1.0 as well as different sizes of the Ligand diameter midpoint box. Xanthine served as the centroid for each grid. Glide Docking^68^ (Schrodinger Inc.) calculations were performed in both Standard Precision (SP) and Extra Precision (XP) modes using the aforementioned parameters. Each run generated over 100 poses.

##### Induced Fit Docking

Induced Fit Docking protocol is a multi-step process designed upon Schrodinger’s Glide and Prime that is capable of accurately predicting non-covalent receptor-ligand complexes. Docking calculations were carried out using the standard protocol of the IFD^68^ tool implemented in Maestro (Schrodinger suite). Initially, Glide docking calculations were performed utilizing a more flexible van der Waals radii scaling, leading to a default maximum of 20 poses per ligand. Subsequently, side-chain prediction and minimization were carried out for each protein-ligand complex using Prime. Finally, redocking of every complex energetically close to the lowest-energy structure was implemented using Glide, and IFDScore (estimation of binding energy) was calculated.

#### MD Simulation System

The protein structures and protein-ligand complexes were constructed using the CHARMM-GUI platform^69^. Each model was embedded into a heterogeneous fully hydrated bilayer measuring 130 Å × 130 Å × 111 Å, composed of YOPC and POPI lipids along with ergosterol in an approximate ratio of 80:80:40. The orientation of the protein within the membrane was achieved by aligning the first principal axis along the Z-axis and subsequently rotating the molecule −90° across the X or Y axis. The system was solvated with TIP3P water molecules and neutralized by adding 150 mM NaCl ions. The assembled simulation system comprised approximately 174,000 atoms, including around 36,000 water molecules. The CHARMM36m^70^ force field was used for both the protein and ligands. Minimization and equilibration were carried out using the default steps provided by CHARMM-GUI to obtain stable structures. The Berendsen thermostat and barostat^71^ were used for equilibration, while the Nose-Hoover thermostat^72^ and Parinello-Rahman barostat^73^ were employed for production. All simulations were conducted using GROMACS software, version 2019.2^74^.

#### Water Maps

Water map calculations were performed on the apo IF, OF, and Occ UapA structures to gain insight into protein solvation and water stability and identify potentially important water molecules. The solvent mapping calculations were performed using the SZMAP algorithm^75^ (OpenEye Inc.) and the following workflow. As a first step, the SPRUCE^76^ tool (OpenEye Inc.) was used for protein structure preparation. Twelve grids were created across the protein, and twelve SZMAP calculations were performed using the -box_mol parameter to achieve faster solvation through the entire pore of UapA. Ligand and solvent molecules from cryoEM were excluded from the calculation. Visualization of the results achieved through VIDA^77^ software (Openeye Inc.)

#### FEP Calculations

Free energy perturbation (FEP) calculations were conducted using the FEP software within Desmond (Schrödinger Inc.)^78^. The IF structure of UapA, along with xanthine and hypoxanthine, were prepared using Schrödinger Maestro software. Hypoxanthine was positioned to align with the experimental CryoEM position of 7H-xanthine in the IF UapA structure. The lipid bilayer was not included in the simulations as it is located away from the xanthine binding site. FEP simulations were performed using default parameters.

## Data availability

The cryo-EM maps of UapA_WT_-Apo-DDM, UapA_WT_-Xan-DDM, UapA_Q408E_-Apo-DDM and UapA_Q408E_-Xan-ND s have been deposited to the Electron Microscopy Data Bank (EMDB) under the accession codes EMD-XXX, EMD-XXX, EMD-XXX and EMD-XXX, respectively. The respective cryo-EM datasets have been deposited to EMPIAR under accession codes EMPIAR-XXX, EMPIAR-XXX, EMPIAR-XXX, and EMPIAR-XXX. The coordinates of the corresponding models have been deposited to the Protein Data Bank (PDB) under accession codes XXX, XXX, XXX, and XXX. The MD input files, starting structures and final structures as representative configurations can be found under https://zenodo.org/records/XXXXXX. Other data are available from the corresponding authors upon request.

