## Supplementary Information for "High-resolution structures of the UapA purine transporter reveal unprecedented aspects of the elevator-type transport mechanism"

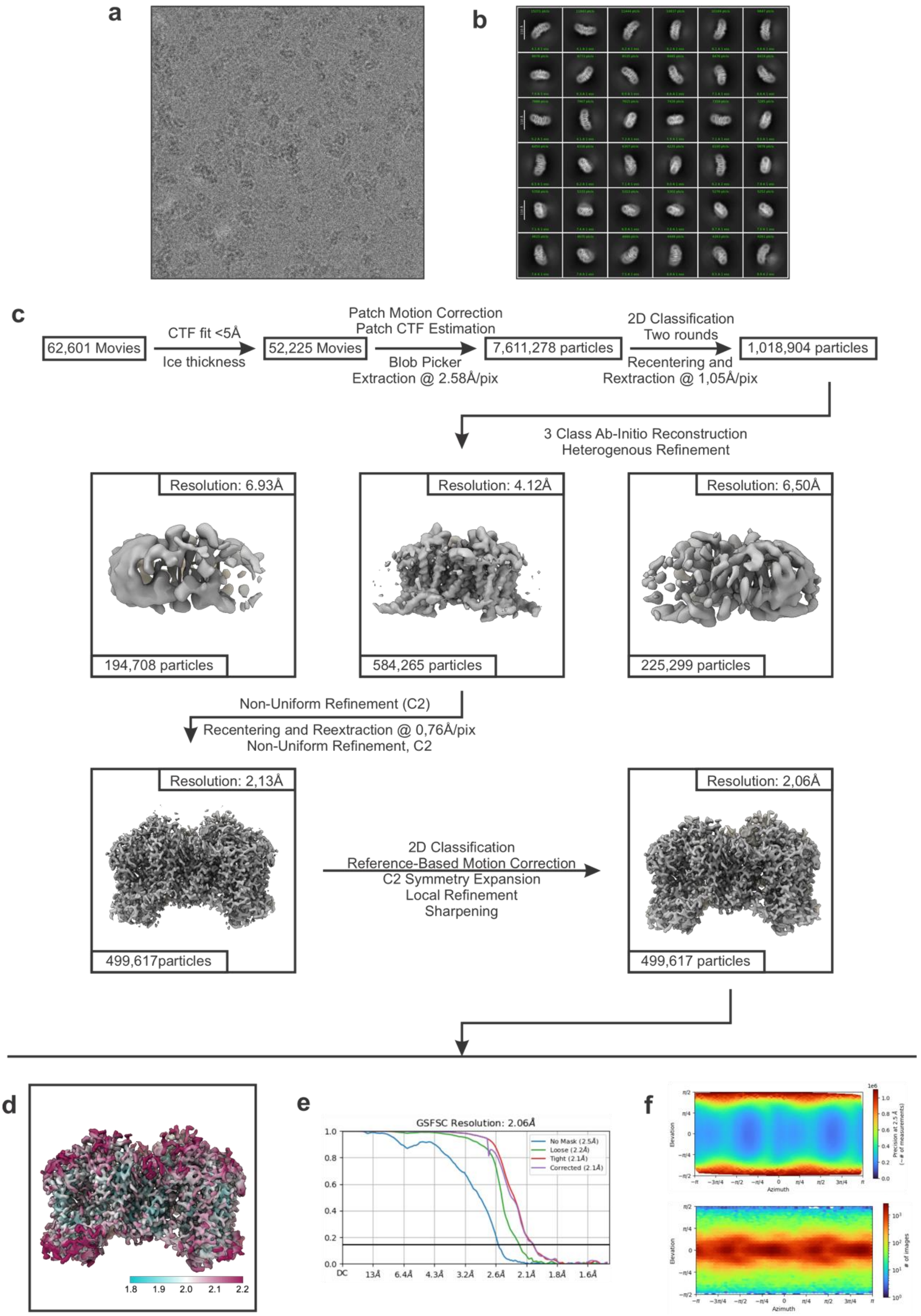


**Supplementary Figure 1: cryoEM analysis of UapA$_{WT}$-Apo-DDM (a-b)** Representative cryoEM micrograph (a) and 2D class averages (b) **(c)** Flowchart for cryo-EM data processing. Please see the Methods sections for more details (**d)** Final cryoEM volume colored according to the local resolution. **(e)** Gold standard FSC **f)** Euler angle distribution.

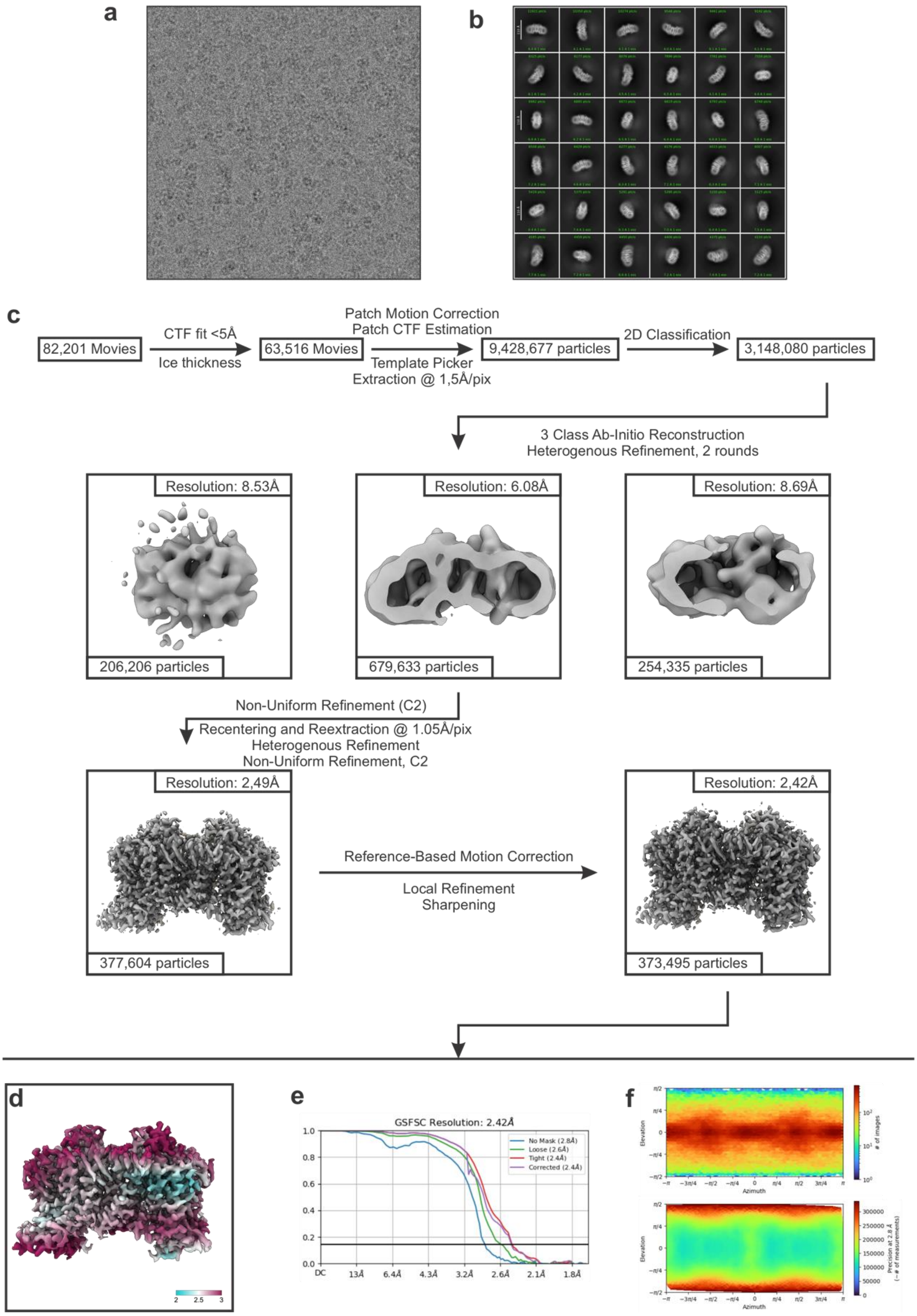


**Supplementary Figure 2: Processing of $UapA_{WT}$-Xan-DDM (a-b)** Representative cryoEM micrograph (a) and 2D class averages (b) **(c)** Flowchart for cryo-EM data processing. Please see the Methods sections for more details **(d)** Final cryoEM volume colored according to the local resolution. **(e)** Gold standard FSC **f)** Euler angle distribution.

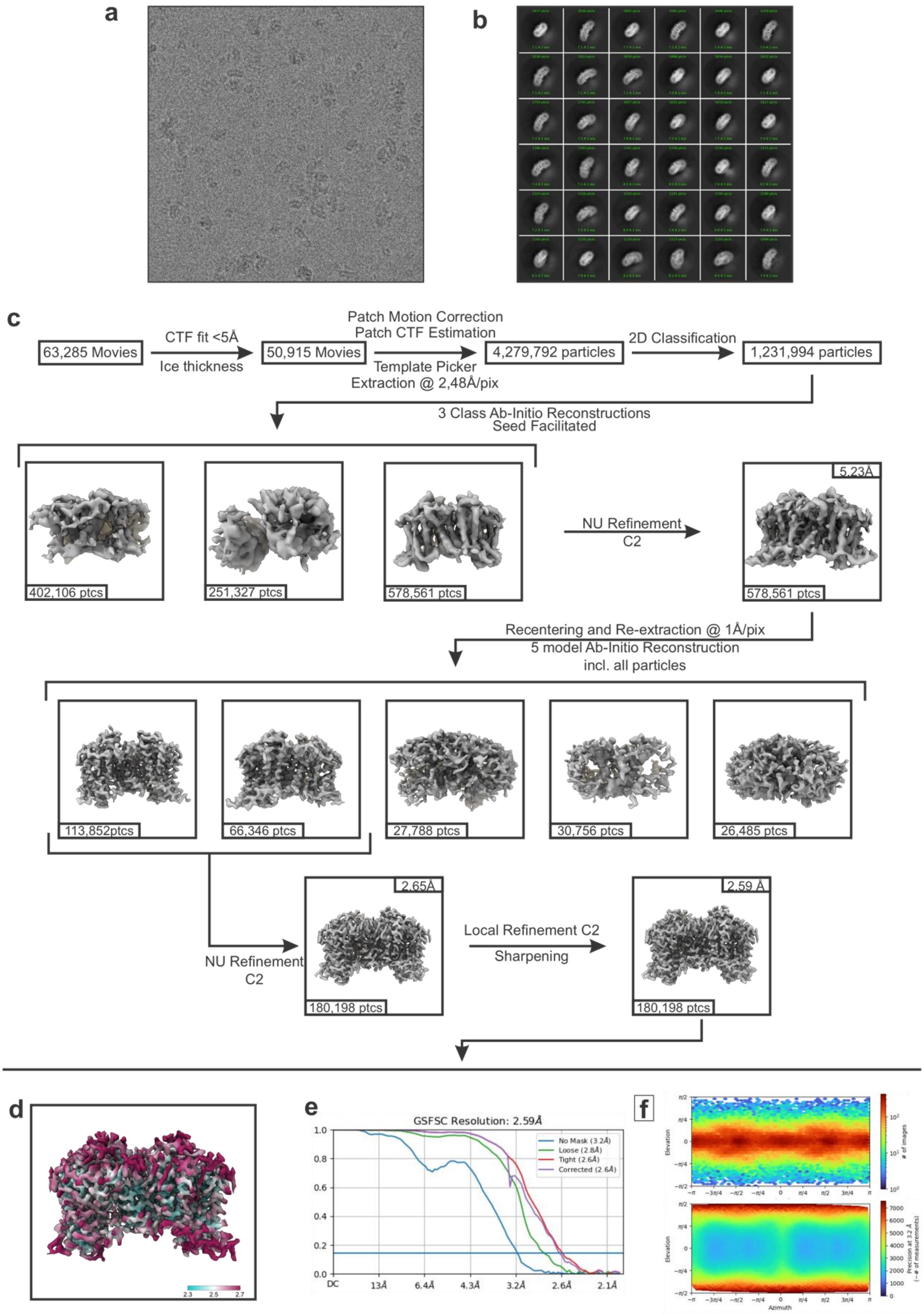


**Supplementary Figure 3: Processing of UapA$_{Q408E}$-Apo-DDM (a-b)** Representative cryoEM micrograph (a) and 2D class averages (b) **(c)** Flowchart for cryo-EM data processing. Please see the Methods sections for more details (**d)** Final cryoEM volume colored according to the local resolution. **(e)** Gold standard FSC **f)** Euler angle distribution.

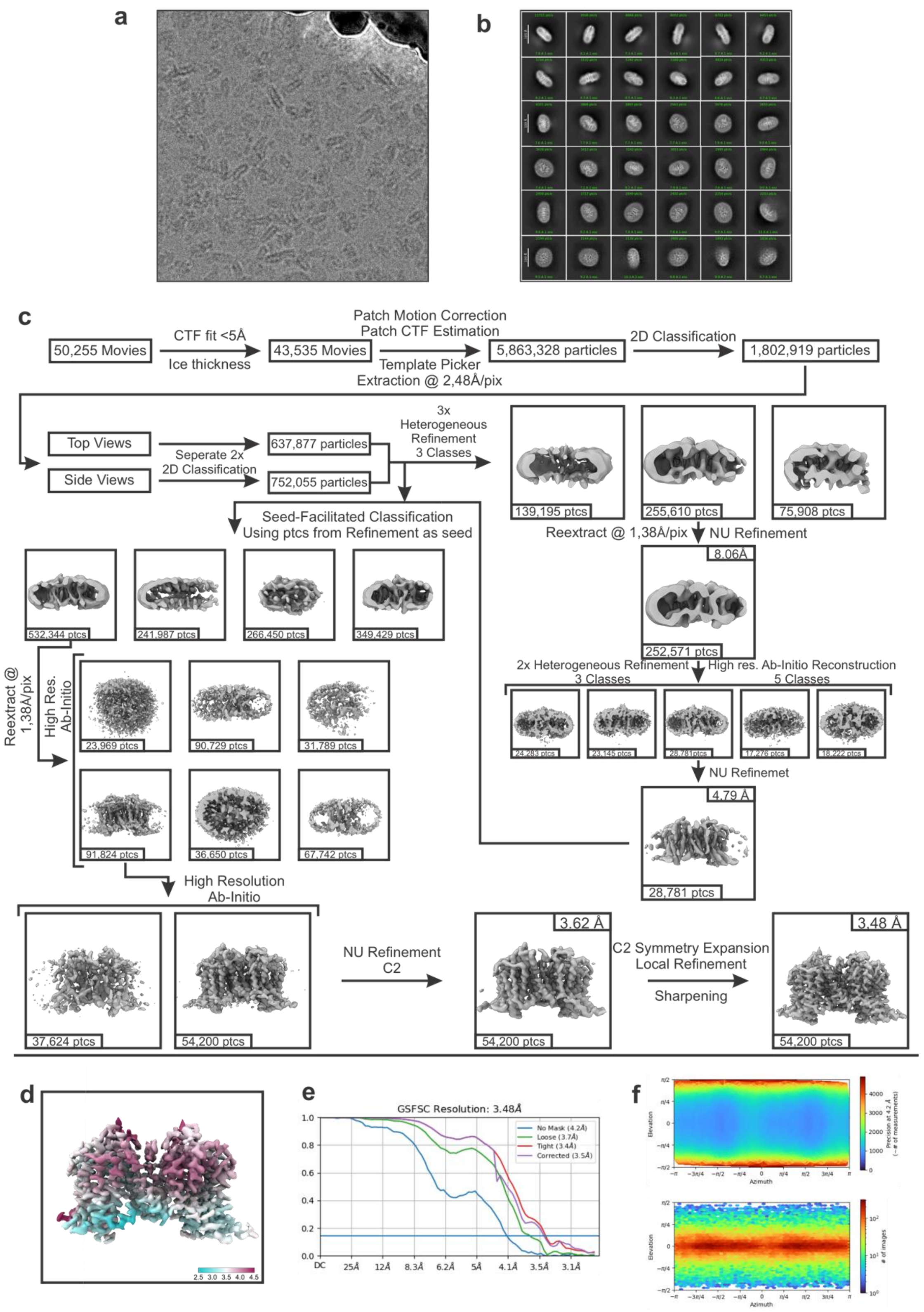


**Supplementary Figure 4: Processing of UapA$_{Q408E}$-Xan-ND (a-b)** Representative cryoEM micrograph (a) and 2D class averages (b) **(c)** Flowchart for cryo-EM data processing. Please see the Methods sections for more details **(d)** Final cryoEM volume colored according to the local resolution. **(e)** Gold standard FSC **f)** Euler angle distribution.

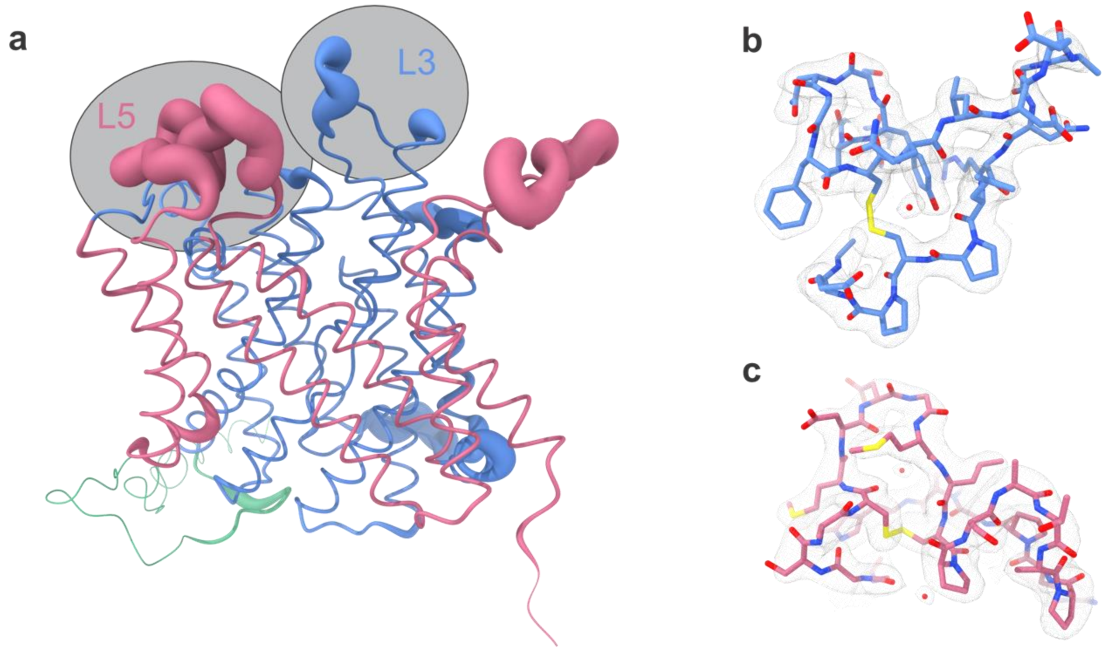


**Supplementary Figure 5: Comparison of the cryoEM structure UapA$_{WT}$-Xan-DDM with crystal structure PDB:5I6C (xanthine-bound UapA$_{G411V}$ in DDM) (a)** RMSD between the two structures, depicted as thickening of the cartoon. **(b)** Closeup of L3, showing the cysteine bond and one water molecule. **(c)** Closeup of L5, showing the novel cystine bond and two water molecules.

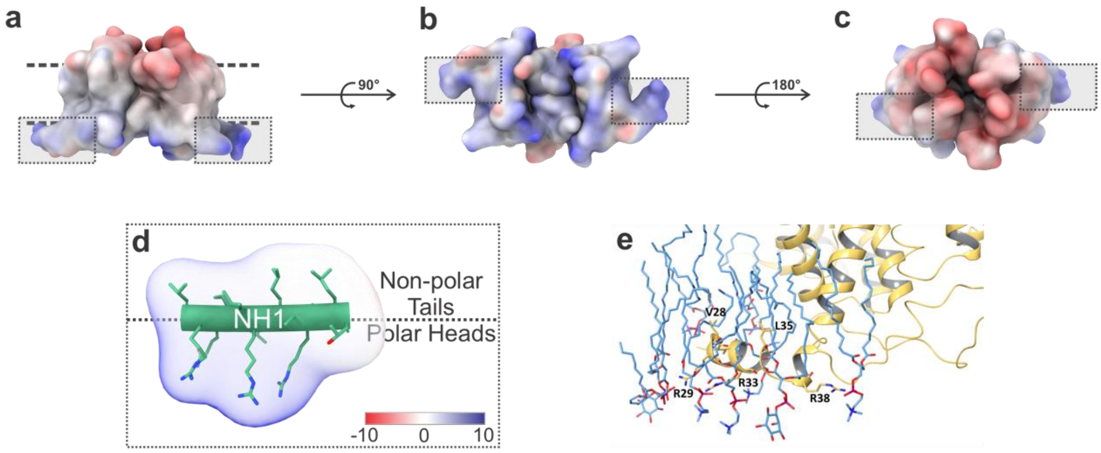


**Supplementary Figure 6: Electrostatic surface of UapA and membrane topology of NH1**
**(a)** Side view **(b)** Bottom view **(c)** Top view **(d)** Close-up of the highly polar NH1, which is perpendicular to and in contact with the membrane. **(e)** Snapshot of MD simulations, showing NH1 immersed in the membrane.

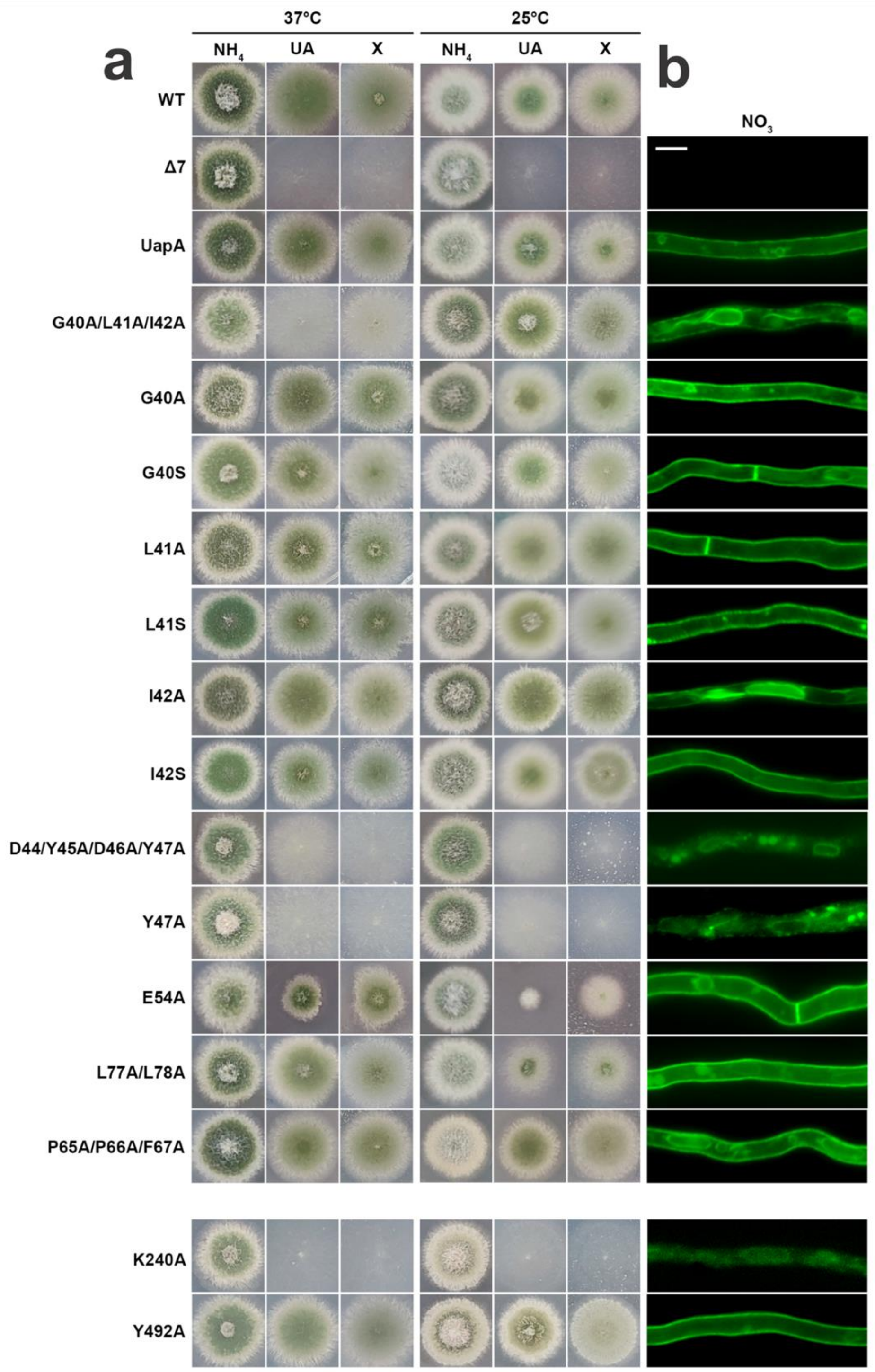


**Supplementary Figure 7: UapA mutations and functional analysis of phenotype (a)** Growth test of UapA mutants. WT is a standard wild-type strains with untagged UapA. Δ7 is a mutant strain genetically lacking all transporters related to nucleobase or nucleoside transport. UapA is a strain expressing wt UapA-GFP in the Δ7 background. All mutants shown are isogenic to UapA and arise from functional single-copy plasmid integration events. In all mutants, UapA is tagged with GFP. Mutants are tested on MM either with a standard nitrogen source unrelated to UapA function ($NH_4$) or on uric acid (UA) or xanthine (X), the main physiological substrates of UapA, as sole nitrogen source, at 25 °C and 37 °C. Positive and negative controls (wt, Δ7, and UapA) are isogenic strains expressing wild-type UapA and a strain lacking all major purine transporters (*uapA*Δ *uapC*Δ *azgA*Δ), respectively. **(b)** Epifluorescence microscopy of UapA mutants.

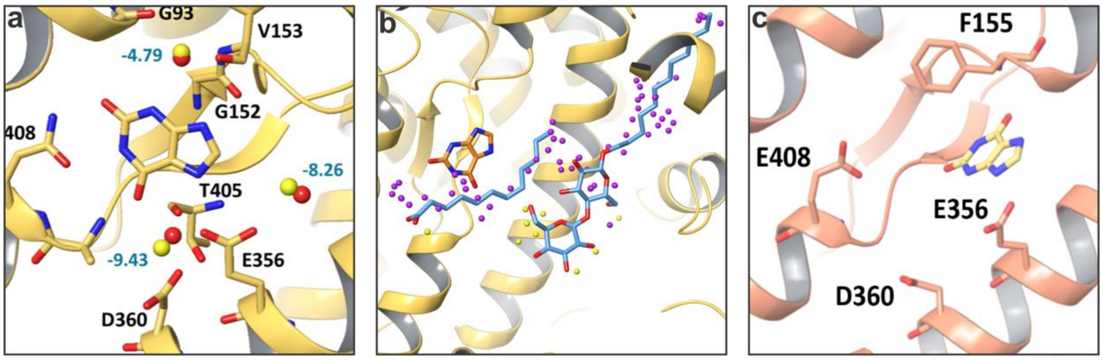


**Supplementary Figure 8: IFD and Solvent Mapping on UapA**

**(a)** SZMAP water prediction. Cryo-EM waters are colored in red, SZMAP in yellow **(b)** High- and low-energy waters predicted around lipid density. Yellow: low-energy waters, Purple: High-energy waters. **(c)** IFD of xanthine in in-silico mutated UapA$_{wt \rightarrow Q408E}$-Xan-DDM is predicted as solved in Nanodisc structure.

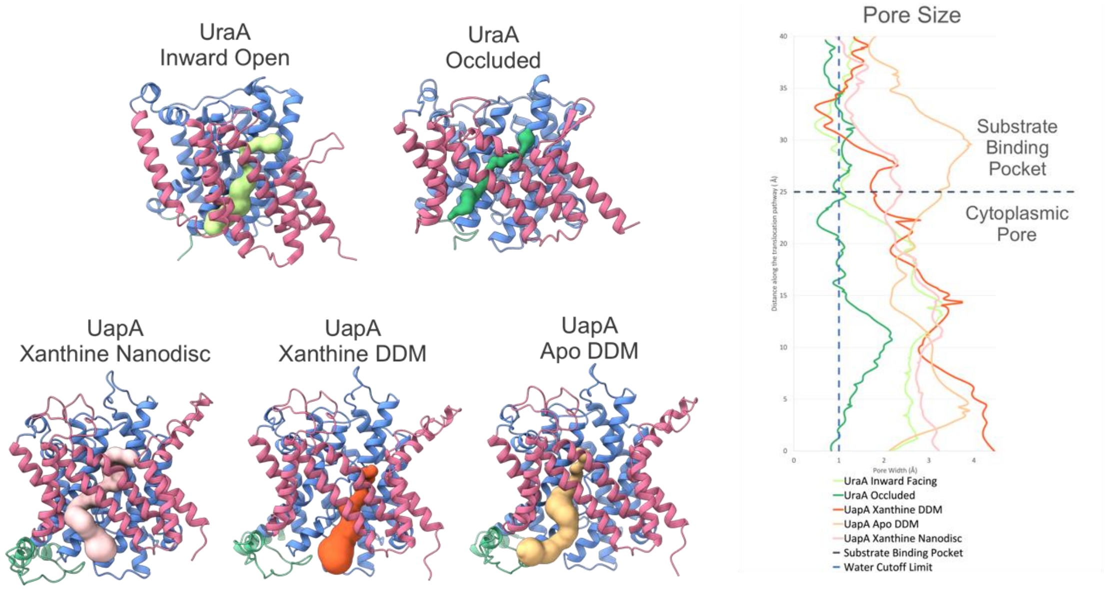


**Supplementary Figure 9: Pore widths of UapA and UraA.** The pores (see skin surfaces) of UraA Inward open (PDB:3QE7) in light green and UraA Occluded (PDB:5XLS) (dark green) are used as conformational controls for the Inward Open and Occluded states and compared with the molecular models of $UapA_{Q408E}$-Xan-ND (pink) $UapA_{WT}$-Xan-DDM (orange) and $UapA_{WT}$-apo-DDM (gold). The plots show the respective pore diameters as a function of distance along the translocation pathway. Here, an occluded state is considered when the pore diameter is less than 1Å below the binding pocket. All structures resolved herein are thus Inward Open.

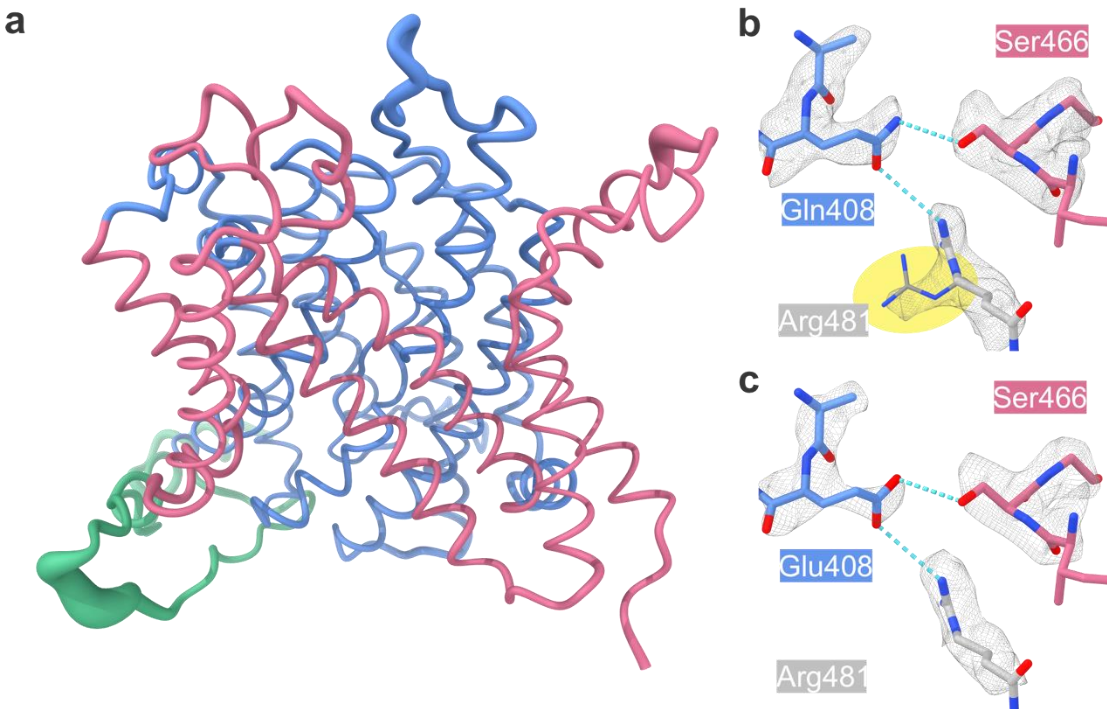


**Supplementary Figure 10: Comparison of UapA$_{WT}$-apo-DDM with UapA$_{Q408E}$-apo-DDM (a)** RMSD between the two structures, depicted as thickening of the cartoon. **(b)** Locking of Gln408 in the WT. Notice the alternative locations for Arg481 in yellow. **(c)** Locking of Glu408 in Q408E.

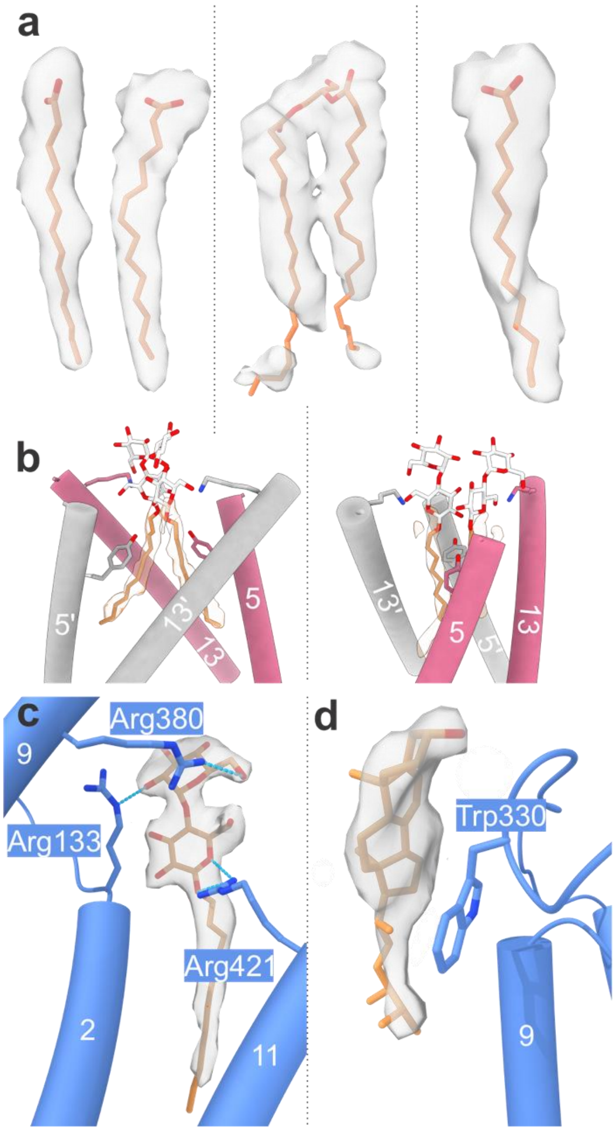


**Supplementary Figure 11: Lipid densities**

**(a)** Typical lipid-like densities, and the fatty acid used to model them. **(b)** Clash of head groups if central lipids are built as DDM molecules **(c)** Putative ergosterol density **(d)** DDM density bound by three arginines.

| Primer name | Sequence |
|---|---|
| UapA-K240-F | GTCTGATTGGAACTGGGTTCGCGGACTGGGCTGGCGGC |
| UapA-K240-R | GCCGCCAGCCCAGTCCGCGAACCCAGTTCCAATCAGAC |
| UapA-Y492-F | GCGTCAATGGCCCTGGGAGCCGGGGCGACGCTGGTG |
| UapA-Y492-R | CACCAGCGTCGCCCCGGCTCCCAGGGCCATTGACGC |
| UapA-G40A-F | CTGACGACCCGCGAGGCCCTGATCGGCGACTATGAC |
| UapA-G40A-R | GTCATAGTCGCCGATCAGGGCCTCGCGGGTCGTCAG |
| UapA-L41A-F | GACGACCCGCGAGGGCGCGATCGGCGACTATGACTATG |
| UapA-L41A-R | CATAGTCATAGTCGCCGATCGCGCCCTCGCGGGTCGTC |
| UapA-I42A-F | GACCCGCGAGGGCCTGGCCGGCGACTATGACTATGGC |
| UapA-I42A-R | GCCATAGTCATAGTCGCCGGCCAGGCCCTCGCGGGTC |
| UapA-G40S-F | CTGACGACCCGCGAGTCCCTGATCGGCGACTATGAC |
| UapA-G40S-R | GTCATAGTCGCCGATCAGGGACTCGCGGGTCGTCAG |
| UapA-L41S-F | GACGACCCGCGAGGGCTCGATCGGCGACTATGACTATG |
| UapA-L41S-R | CATAGTCATAGTCGCCGATCGAGCCCTCGCGGGTCGTC |
| UapA-I42S-F | GACCCGCGAGGGCCTGTCCGGCGACTATGACTATGGC |
| UapA-I42S-R | GCCATAGTCATAGTCGCCGGACAGGCCCTCGCGGGTC |
| UapA-E54A-F | CTTCCTTTTCCGGCCCGCGCTACCCTTCATGAAGAAAG |
| UapA-E54A-R | CTTTCTTCATGAAGGGTAGCGCGGGCCGGAAAAGGAAG |
| UapA-L77A/L78A-F | CTCAACGAGAAGATTCCCGTGGCCGCGGCGTTTATCCTGGGTCTTC |
| UapA-L77A/L78A-R | GAAGACCCAGGATAAACGCCGCGGCCACGGGAATCTTCTCGTTGAG |
| UapA-P65A/P66A/F67A-F | GAAGAAAGATCCACGGGCGGCCGCAGCTTTTGGCCTCAACGAGAAG |
| UapA-P65A/P66A/F67A-R | CTTCTCGTTGAGGCCAAAAGCTGCGGCCGCCCGTGGATCTTTCTTC |

**Supplementary table 1.** Primers used for site-directed mutagenesis. F denotes forward and R reverse. Sequences are written 5’ to 3’.
